# A chromatin relay from AIRE to ETS transcription factors sustains peripheral antigen expression in the thymic mimetic cells to ensure central tolerance

**DOI:** 10.64898/2026.02.03.703355

**Authors:** Kenta Horie, Takahisa Miyao, Yuko Yamagata, Yuki Takakura, Fuyoko Suzuki, Naho Hagiwara, Rin Endo, Maki Miyauchi, Hiroto Ishii, Wataru Muramatsu, Yuki Goda, Tatsuya Ishikawa, Jun-ichiro Inoue, Nobuaki Yoshida, Noritaka Yamaguchi, Georg A. Hollander, Eiryo Kawakami, Shigeo Murata, Nobuko Akiyama, Taishin Akiyama

## Abstract

Medullary thymic epithelial cells (mTECs) establish central tolerance by expressing a diverse repertoire of peripheral tissue-specific antigens (TSAs). This diversity is regulated not only by the transcriptional regulator AIRE, but also by lineage-defining factors in tissue-mimetic, AIRE-negative descendants of Aire⁺ mTECs (post-Aire mimetic TECs). However, whether and how prior AIRE activity contributes to TSA expression in post-AIRE mimetic TECs remains unclear. Here, we identify the ETS transcription factors, EHF and ELF3, as key regulators that sustain AIRE-primed gene expression in these cells. EHF and ELF3 preferentially bind distal genomic regions which are rendered accessible in advance by AIRE. Combined loss of EHF and ELF3 disrupts expression of AIRE-regulated genes expressed in mimetic TECs, leading to tissue-selective autoimmunity. Our findings reveal a relay mechanism in which AIRE primes chromatin for ETS factors to maintain TSA expression in post-AIRE mimetic TECs, thereby safeguarding self-tolerance.

## Introduction

Medullary thymic epithelial cells (mTECs) are essential for establishing T cell tolerance and preventing autoimmune disease^1, 2^. By presenting a diverse array of peripheral tissue–specific antigens (TSAs) to developing thymocytes, mTECs eliminate autoreactive T cells or divert them into the regulatory T cell (Treg) lineage^1, 2^. A central regulator of this process is the transcriptional regulator AIRE, which drives the expression of a subset of TSAs in mTECs^3^, and loss-of-function mutations in AIRE cause the human autoimmune syndrome autoimmune polyendocrinopathy-candidiasis-ectodermal dystrophy (APECED)^4^. In the postnatal thymus, mTECs are heterogeneous and can be distinguished by the expression of surface markers and functional genes, including AIRE and the chemokine CCL21^5^. mTECs expressing high levels of MHC class II and AIRE (Aire^+^ mTECs) arise from mTEC progenitors and subsequently differentiate into post-Aire mimetic TECs, which lose AIRE expression.

Aire⁺ and post-Aire mimetic TECs are chiefly responsible for TSA expression through mechanistically distinct transcriptional programs. In Aire⁺ mTECs, the transcriptional regulator AIRE drives TSA expression in a largely stochastic manner^6^. In contrast, post-Aire mimetic TECs downregulate AIRE and maintain TSA expression in a more coordinated, lineage-restricted fashion by adopting transcriptional programs that mimic various cell lineages, including tuft, endocrine, and microfold cells^7^. Thus, post-Aire mimetic TECs retain a core thymic identity while acquiring a “mimetic” property through the activation of lineage-specific gene expression. However, because post-Aire mimetic TECs arise from Aire⁺ mTECs^8, 9, 10, 11, 12^, it is possible that TSA expression in these cells remains influenced by prior AIRE activity, despite the loss of AIRE expression. Consistent with this idea, the previous study has shown that the Aire deficiency altered transcriptional profiles of Aire-negative thymic tuft cells^13^. However, the mechanistic link between AIRE function and the establishment of TSA programs in post-Aire mimetic TECs has remained unresolved.

The E26 transformation-specific (ETS) transcription factor family consists of approximately 28 members in mammals, characterized by a conserved DNA-binding domain that recognizes GGAA/T motifs^14^. These factors regulate a wide range of biological and pathological processes^15^, including development, differentiation, immune function, and oncogenesis. Earlier work using promoter assays in cell culture suggested that ETS proteins contribute to the regulation of Aire expression^16^. In addition, the ETS family member Spi-B has been shown to control mTEC gene expression^7, 17^ and co-stimulatory molecule surface expression^17^. More recently, analyses combining ATAC-seq footprinting and ChIPmentation suggested ETS subfamilies, including E74 Like ETS transcription factor (ELF), epithelium-specific ETS transcription factor, ETS2 repressor factor, and polyomavirus enhancer activator 3, as potential regulators of mTEC gene expression in late differentiation stages of mTECs^18^. However, whether and how ETS family members functionally intersect with AIRE to regulate mTEC biology has not been addressed.

Here, we describe an interplay between AIRE and the ETS factors, ETS homologous factor (EHF) and E74 Like ETS transcription factor 3 (ELF3), in sustaining gene expression in post-Aire mimetic TECs. In Aire⁺ mTECs, AIRE establishes chromatin footprints without fully opening these sites. Upon differentiation into post-Aire mimetic TECs, EHF and ELF3 recognize these “AIRE-primed” regions and activate peripheral antigen expression, thereby reinforcing central tolerance.

## Results

### AIRE priming contributes to gene expression in post-Aire mimetic TECs

Because post-Aire mimetic TECs arise from Aire⁺ mTECs, AIRE activity can influence gene expression programs of post-Aire mimetic TECs after its expression has ceased. To address this, we analyzed previously reported scRNA-seq data from thymic epithelial cells (TECs) of control and Aire-deficient (*Aire*⁻^/^⁻) mice (https://doi.org/10.1101/2025.08.06.668799). We focused on mature mTECs, Aire^+^ mTECs, and post-Aire mimetic TECs, by excluding Ccl21⁺ mTECs, cortical TECs (cTECs), transit-amplifying TECs (TA-TECs)^19, 20, 21^, and progenitor populations^22, 23^ (Supplementary Figure S1). Subclustering analysis on mature mTECs resolved 10 distinct clusters, with assignment: Aire^+^ mTEC (cluster 0, 2, and 6), Late Aire mTECs (cluster 1) and post-Aire mimetic TECs including tuft-like mTECs (cluster 3, 4, 5, 7, 8, and 9) (Figure 1A), providing a framework to examine how AIRE-dependent transcriptional programs evolve during the transition from Aire⁺ to post-Aire states.

**Figure 1.**
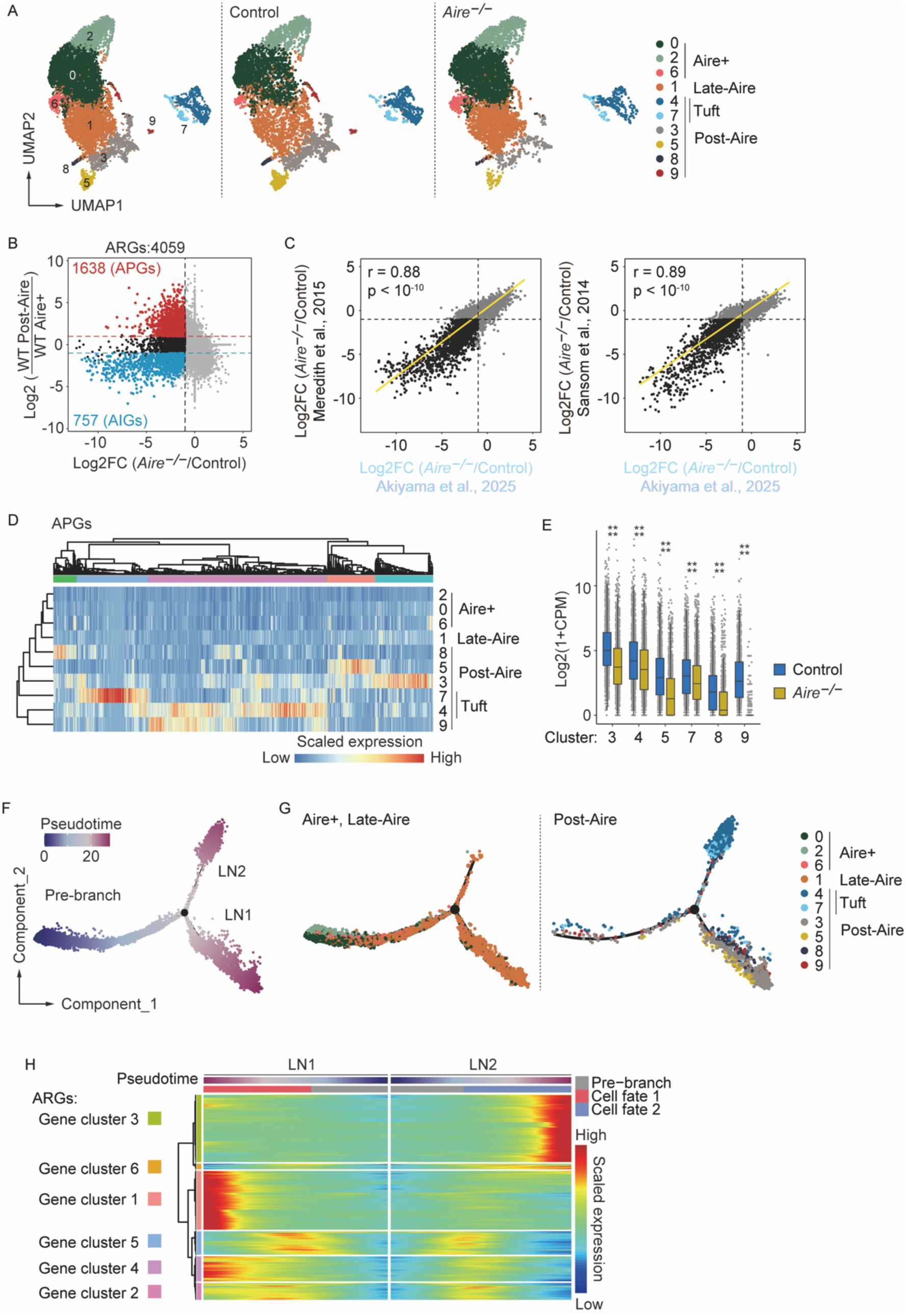
Aire depletion affects gene expression programs in post-Aire mimetic TECs. A. UMAP of Aire^+^ mTECs and post-Aire mimetic TECs extracted from whole TEC scRNA-seq shown in supplementary data 1, resulting from dimension reduction and clustering analysis. B. FC/FC plots of genes comparing fold changes of control post-Aire mimetic TECs versus control Aire^+^ mTECs and *Aire*^−/–^ versus control of all mature mTECs. Black dots indicate significantly down-regulated genes by *Aire*^−/–^ compared to control (Aire-regulated genes: ARGs). Red dots indicate significantly higher ARGs in post-Aire mimetic TECs compared to Aire^+^ mTECs (Aire-primed genes: APGs). Blue dots indicate significantly lower ARGs by Aire^+^ mTECs compared to post-Aire mimetic TECs (Aire-induced genes: AIGs). Significantly altered genes were defined by absolute Log2FC > 1 and FDR < 0.05. C. FC/FC plots between *Aire*^−/–^ and control comparing mTEC^hi^ bulk RNA-seq from the previous studies versus pseudobulk mature mTECs from our scRNA-seq. R- and p-values were calculated using Pearson correlation analysis. D. A heatmap showing normalized APG expression. E. Box plots showing log2-transformed CPM of APGs from Figure. 1B in indicated cell clusters. P-values were calculated using Wilcoxon rank-sum test with Benjamini-Hochberg correction. F. Component graphs show the results of DDRTree dimensionality reduction analysis applied to single-cell RNA sequencing data shown in Figure. 1A. A black dot indicates the branching point. Pseudotime is projected on component graphs. G. Component graphs show the results of DDRTree dimensionality reduction analysis applied to single-cell RNA sequencing data shown in Figure. 1A. A black dot indicates the branching point. Clusters are projected on component graphs. H. A heatmap showing scaled ARG expression along the pseudotime trajectory for each lineage. Genes are clustered based on the similarity of gene expression patterns along the pseudotime trajectory. LN1: Lineage 1; LN2: Lineage2

As expected, comparison of *Aire*⁻^/^⁻ and control total mature mTECs, encompassing all mature mTEC clusters identified in the scRNA-seq analysis, revealed broad transcript downregulation (4,059 genes; Figure 1B). Given that the Aire-dependent expression changes we observed closely matched those reported in prior datasets^6, 24^ (Figure 1C), we designate these downregulated genes as Aire-regulated genes (ARGs) for subsequent analyses (Table 1). Despite the downregulation of AIRE during differentiation, 1,638 ARGs displayed higher expression in post-Aire mimetic TECs than in Aire⁺ mTECs (red dots in Figure 1B), and were therefore classified as Aire-primed genes (APGs). By contrast, 757 ARGs were more highly expressed in Aire⁺ mTECs and were classified as Aire-induced genes (AIGs; blue dots in Figure 1B). Examination of individual post-Aire clusters indicated that these APGs were not uniformly expressed but appeared more prominent in specific clusters (Figure 1D). Importantly, their expression remained dependent on prior AIRE activity (Figure 1E), highlighting a cluster-selective persistence of the AIRE-dependent transcription program in post-Aire mimetic TECs.

To further resolve the relation between this differentiation process and Aire-regulated gene expression, we next performed pseudotime trajectory analysis using the DDRTree algorithm implemented in Monocle 2^25^ (Figure 1F). Clusters 0, 2, and 6, composed primarily of Aire⁺ mTECs, localized upstream of the trajectory branching point (Figure 1G), consistent with their identity as early differentiation stages. Late-Aire mTECs (cluster 1) were distributed more broadly along the pseudotime axis, suggesting transitional states spanning pre- and post-branch phases. In contrast, post-Aire mimetic TECs were positioned predominantly downstream of the branching point, indicative of more differentiated states (Figure 1F). Notably, the post-branch trajectory bifurcated into two distinct lineages. One branch (lineage 1, LN1) consisted of conventional (non-tuft) post-Aire clusters together with Late-Aire mTECs, whereas the other branch (lineage 2, LN2) was composed mainly of tuft-like TEC clusters (Figure 1E and F). This bifurcation suggests that post-Aire mimetic TECs diverge into two major lineage programs, with LN1 representing conventional post-Aire mimetic TECs and LN2 adopting a tuft-like TEC fate.

To investigate dynamic changes in ARG expression along pseudotime, we applied branched expression analysis modeling (BEAM) to ARGs^25^, which identified five distinct ARG clusters (Figure 1H, Table 2). ARGs in cluster 1 and cluster 4 were upregulated toward the end of the LN1 trajectory, while those in cluster 3 were upregulated at the end of LN2 in an Aire-dependent manner. In contrast, clusters 2 and 5 exhibited transient upregulation earlier along the trajectory. Thus, some subsets of ARGs showed low expression in Aire⁺ mTECs but became highly expressed during the differentiation into post-AIRE mimetic TECs.

Overall, these findings reveal two distinct modes of AIRE regulation. First, AIRE directly induces the expression of a subset of ARGs in Aire⁺ mTECs. Second, although AIRE expression is downregulated upon differentiation into post-Aire mimetic TECs, prior AIRE activity enables a distinct subset of ARGs to reach maximal expression only after this transition. Thus, AIRE functions not only as a direct transcriptional activator in Aire⁺ mTECs but also as a priming factor that establishes a transcriptional program subsequently amplified in post-Aire mimetic TECs.

### Dual roles of AIRE in chromatin regulation during the transition to post-Aire mimetic TECs

Given AIRE’s role in priming ARG expression in post-Aire mimetic TECs, we next examined its impact on chromatin accessibility using scATAC-seq data of *Aire*^−/–^ and control TECs from the previous study (https://doi.org/10.1101/2025.08.06.668799). To link between chromatin accessibility and gene expression, we integrated these scATAC-seq data with single-cell RNA+ATAC multiome profiles from the other previous study^26^. After removing low-quality cells, we retained 31,501 cells in total: 16,065 control, 6,761 *Aire*^−/–^, and 8,675 control wild-type multiome cells. Dimensionality reduction and clustering analysis of the combined accessibility matrix resolved 20 clusters based on chromatin accessibility profiles (Supplementary Figure S2). On the basis of canonical marker gene expression, we then extracted Aire^+^ and post-Aire mimetic TECs and re-clustered these subsets for downstream analyses, yielding 11 distinct clusters (Figure 2A). As expected, numerous genomic regions exhibited altered chromatin accessibility during the transition from Aire⁺ mTECs to post-Aire mimetic TECs (Figure 2B). In post-Aire mimetic TECs, 6200 loci were significantly less accessible in *Aire*⁻^/^⁻ mice (Figure 2C), indicating that prior AIRE activity is required to establish or maintain accessibility at these regions. Integration of our scATAC-seq data with published AIRE ChIP-seq profiles^27^ further revealed that the loci closed in *Aire*⁻^/^⁻ post-Aire mimetic TECs were preferentially bound by AIRE in Aire⁺ mTECs (Figure 2D). These findings support a model in which AIRE primes specific genomic sites during the Aire⁺ mTEC stage, thereby enabling their accessibility to persist in post-Aire mimetic TECs despite the loss of AIRE expression.

**Figure 2.**
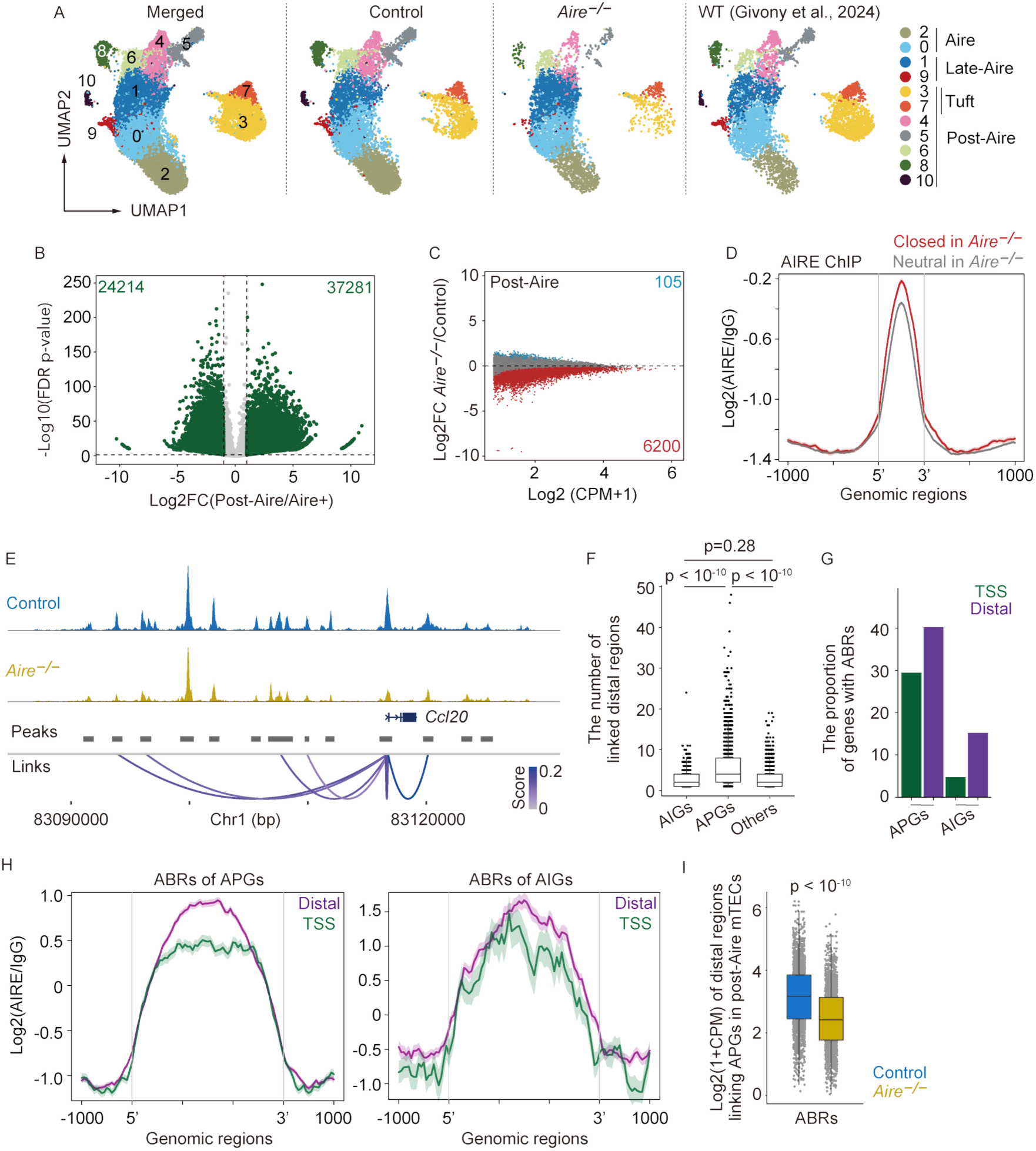
AIRE pre-binds some distal regulatory elements that subsequently gain chromatin accessibility in post-Aire mimetic TECs. A. UMAP of Aire^+^ mTECs and post-Aire mimetic TECs extracted from whole TEC scATAC-seq shown in supplementary data 2, resulting from dimension reduction and clustering analysis. B. A volcano plot indicating differentially accessible regions in post-Aire mimetic TECs compared to Aire^+^ mTECs from Control. Each gray dot represents a peak, and each green dot represents significantly altered accessible region which are defined as FDR < 0.05 and absolute Log2FC > 1. C. A MA plot of log-transformed CPM of Control and log2 fold changes of chromatin accessibility between *Aire*^−/–^ and Control in post-Aire mimetic TECs. Each dot represents a peak significantly opened in post-Aire mimetic TECs compared to Aire+ mTECs from Control, defined in Figure 2B. Each red dot represents a significantly closed peak, and each blue dot represents a significantly opened peak by *Aire*^−/–^ compared to Control in post-Aire mimetic TECs, which are defined as FDR < 0.05. D. A profile of AIRE ChIP-seq (Bansal K et al., 2017) at Aire-closed regions and Aire neutral regions in post-Aire mimetic TECs which were defined as red or gray dots in Figure 2C, respectively. E. Peak tracks of representative ARGs significantly down-regulated by Aire^−/–^ compared to Control. Arch plots link peaks correlated with gene expression. Scores mean Z-score calculated by using the correlation coefficient. F. Box plots indicating the number of distal regions significantly correlated with each ARG expression. Each gray dot represents an ARG, and ARGs were divided into three groups (AIGs, APGs, or others) based on expression patterns in Figure 1B. P-values were calculated using Wilcoxon rank-sum test with Benjamini-Hochberg correction. G. Bar plots indicating the proportions of ARGs with ABRs. H. Profiles of AIRE ChIP-seq (Bansal K et al., 2017) at Aire-bound distal regions or transcription start sites (TSS) linked with APG expression (left) or AIG expression (right). I. Box plots indicating log-transformed read counts at distal regions linking APG expression in post-Aire mimetic TECs. Distal regions were divided into Aire-bound regions (ABRs) and non-ABRs based on AIRE ChIP-seq (Bansal K et al., 2017). P-value was calculated using Wilcoxon rank-sum test with Benjamini-Hochberg correction.

AIRE binds both distal regulatory elements, such as enhancers, and transcription start sites (TSSs)^26,44^. To investigate how AIRE-dependent transcriptional programs are maintained in post-Aire mimetic TECs, we identified distal genomic regions whose chromatin accessibility was significantly correlated with gene expression using scEpiMultiome data of wild-type mTECs^25^ (Figure 2E). By comparing these distal genomic regions related to two classes of ARGs defined from the scRNA-seq analysis (Figure 1; AIGs, preferentially expressed in Aire⁺ mTECs, and APGs, preferentially upregulated in post-Aire mimetic TECs), we revealed that APGs were associated with a greater number of distal regulatory elements, such as enhancers located far from TSSs, compared with AIGs (Figure 2F). Consistently, APGs contained more AIRE-bound sites within their regulatory landscapes than AIGs, both at promoters and distal regions (Figure 2G), with AIRE occupancy particularly enriched at distal elements (Figure 2H). Notably, the accessibility of these distal AIRE-bound regions was significantly reduced in Aire⁻^/^⁻ cells (Figure 2I). Together, these findings suggest that AIRE establishes accessibility at distal enhancers of APGs, thereby priming them for transcriptional activation after AIRE expression has ceased. Thus, AIRE may exert dual functions in Aire⁺ mTECs: while compacting chromatin at some loci, it simultaneously primes distal regulatory elements that ensure sustained expression of a distinct gene set in post-Aire mimetic TECs.

### Identification of EHF and ELF3 as regulators of AIRE-primed gene expression in post-Aire mimetic TECs

We next investigated the regulatory mechanisms that drive APG expression in post-Aire mimetic TECs. To this end, we performed motif enrichment analysis on distal regulatory regions identified as AIRE-primed in our scATAC-seq dataset and published AIRE ChIP-seq profiles^27^ (Figure 2E-I). We first selected motifs that were present in more than 50% of peaks within these regions, ranked among the top 100 by statistical significance, and exhibited more than a two-fold enrichment score (Figure 3A). We then incorporated scRNA-seq data to filter for transcription factors with a mean expression greater than 1 in post-Aire mimetic TECs and exhibiting more than fourfold upregulation relative to Aire⁺ mTECs (Figure 3B). This integrated approach identified the ETS family members ELF3 and EHF as candidate regulators binding to Aire-primed distal regulatory elements in post-Aire mimetic TECs. These factors share highly similar binding motifs (Figure 3C) and show nearly identical expression patterns across post-Aire mTEC subsets, with the exception of lower Ehf expression in cluster 8 (Figure 3D), suggesting functional redundancy. Notably, ELF3- and EHF-associated motif activity was reduced in *Aire*⁻^/^⁻ post-Aire mimetic TECs (Figure 3E), supporting a model in which AIRE primes distal enhancers that are subsequently engaged by ELF3 and EHF to drive APG expression.

**Figure 3.**
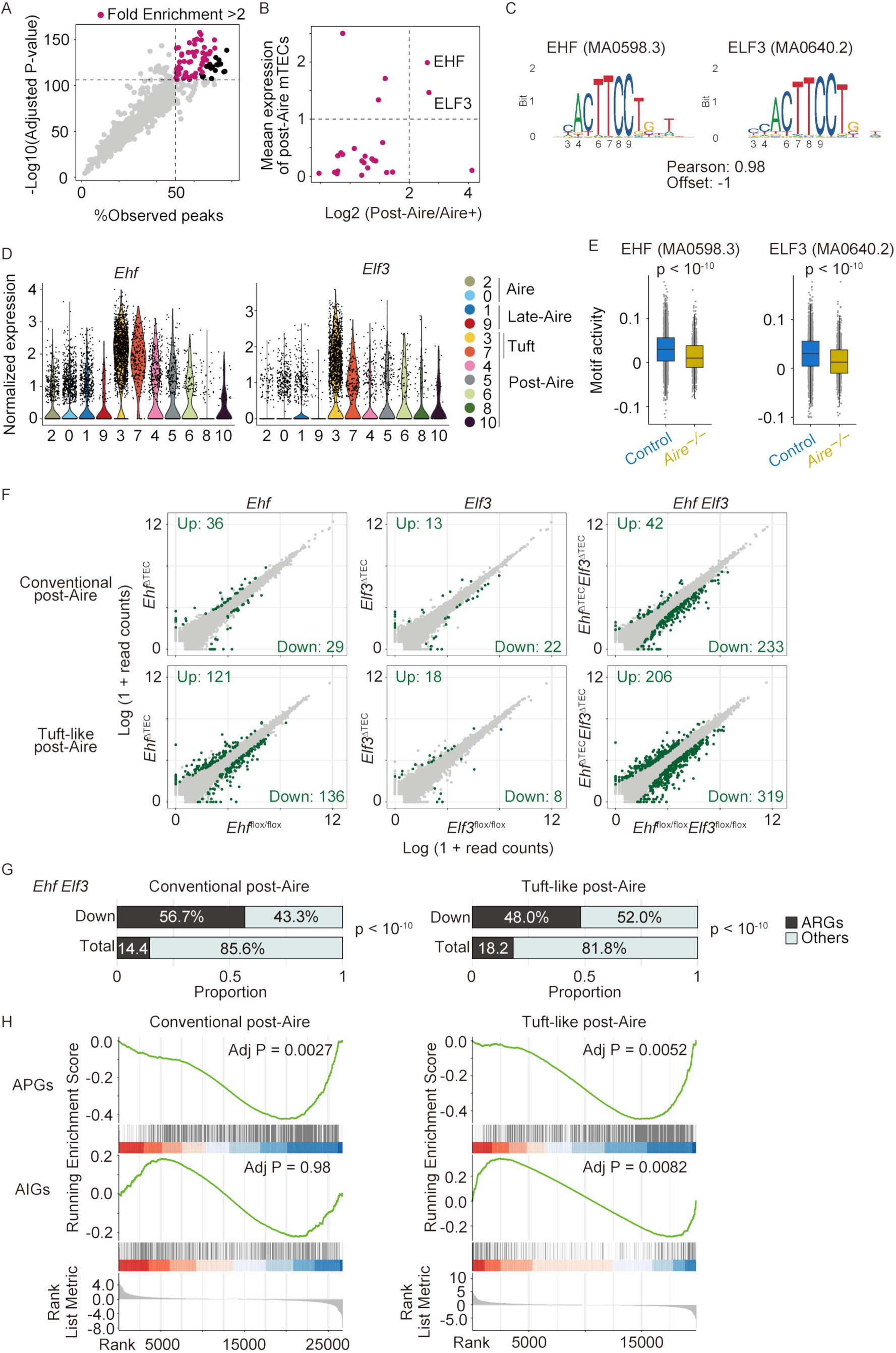
EHF and ELF3 regulate gene expression in post-Aire mimetic TECs, with binding motifs enriched at AIRE-primed genes. A. Scatter plots showing enriched motifs of transcription factor binding DNA sequences at differentially accessible Aire-bound distal regions linking APG expression in post-Aire mimetic TECs. The x-axis indicates the percentage of peaks with a target motif in the total number of peaks, and the y-axis indicates log10-transformed p-values adjusted with Benjamini-Hochberg correction. Each dot represents a transcription factor binding motif. Black dots represent motifs with a top100 minimum p-value or an % observed peak over 50, and magenta dots denote transcription factor motifs with fold enrichment scores exceeding 2. B. A scatter plot showing transcription factors meeting both criteria, described above (Figure 3A). The x-axis indicates log2 fold changes between post-Aire mimetic TECs and Aire^+^ mTECs in control, and the y-axis indicates mean expression in post-Aire mimetic TECs from the scRNA-seq data (Figure 1). C. Motif sequences of EHF (MA0598.3) and ELF3 (MA0640.2) form JASPAR database. D. Violin plots indicating gene expression of Ehf and Elf3 in indicated clusters. E. Violin plots indicating the motif activity of EHF (MA0598.3) and ELF3 (MA0640.2) in post-Aire mimetic TECs. P-values were calculated using Wilcoxon rank-sum test with Benjamini-Hochberg correction. F. Scatter plots comparing log-normalized gene expression values of TEC subpopulations in indicated genotypes. Each dot represents a gene, and green plots indicates significantly altered genes, defined by absolute fold change > 2 and FDR < 0.05. Each genotype has three biological replicates (N = 3). G. Bar plots comparing proportions of ARGs in indicated mTEC subpopulations. P-values were calculated using the chai-squared test. H. GSEA analysis of APGs and AIGs on differential gene expression profiles of conventional post-Aire mimetic TECs and tuft-like post-Aire mimetic TECs between *Ehf*^flox/flox^ *Elf3*^flox/flox^ and *Ehf*^ΔTEC^*Elf3*^ΔTEC^.

To test the roles of these factors on gene expression of post-Aire mimetic TECs, mice deficient in *Ehf* or *Elf3* in TECs by using Foxn1-Cre (*Ehf*^ΔTEC^ and *Elf3*^ΔTEC^) were established. Considering the functional redundancy between EHF and ELF3, we established mice deficient in both Ehf and Elf3 in TECs (*Ehf*^ΔTEC^*Elf3*^ΔTEC^). Flow cytometric analysis of TECs using several surface markers showed no significant influences of these deficiencies in the development of mTEC subsets and cTECs (Supplementary Figure S3).

We performed bulk RNA-seq analysis of sorted mTEC subsets and cTECs from mutant mice (Figure 3F and Supplementary Figure S4). Differential expression analysis showed that deletion of *Ehf* had a mild effect on gene expression in tuft-like cells, whereas deletion of *Elf3* alone had minimal impact across TEC subsets. In contrast, combined deletion of *Ehf* and *Elf3* caused extensive transcriptional changes in both conventional post-Aire and tuft-like mTECs (Figure 3F), indicating that these transcription factors function redundantly to regulate gene expression in post-Aire cells. Notably, downregulated genes in double-mutant conventional post-Aire and tuft-like mTECs included approximately 50% of ARGs (Figure 3G), a proportion significantly higher than expected based on their frequency among all detected genes. Moreover, APGs were preferentially downregulated in both conventional post-Aire and tuft-like mTECs (Figure 3H), while AIGs were not significantly enriched in the down-regulated genes by the double deficiencies. These findings suggest that EHF and ELF3 cooperatively sustain the expression of a subset of AIRE-primed genes in post-Aire mimetic TECs after AIRE expression has ceased.

We next investigated the functional convergence between AIRE and EHF/ELF3 in post-AIRE mimetic TECs. Bulk RNA-seq analysis of sorted tuft-like mTECs revealed that gene expression fold-change values relative to control were positively correlated between *Ehf^Δ^*^TEC^*Elf3*^ΔTEC^ and *Aire*⁻^/^⁻ mice for APGs, but not for AIGs (Figure 4A). These results indicate that AIRE and EHF/ELF3 converge functionally on the regulation of APGs, despite acting at distinct stages of mTEC differentiation.

**Figure 4.**
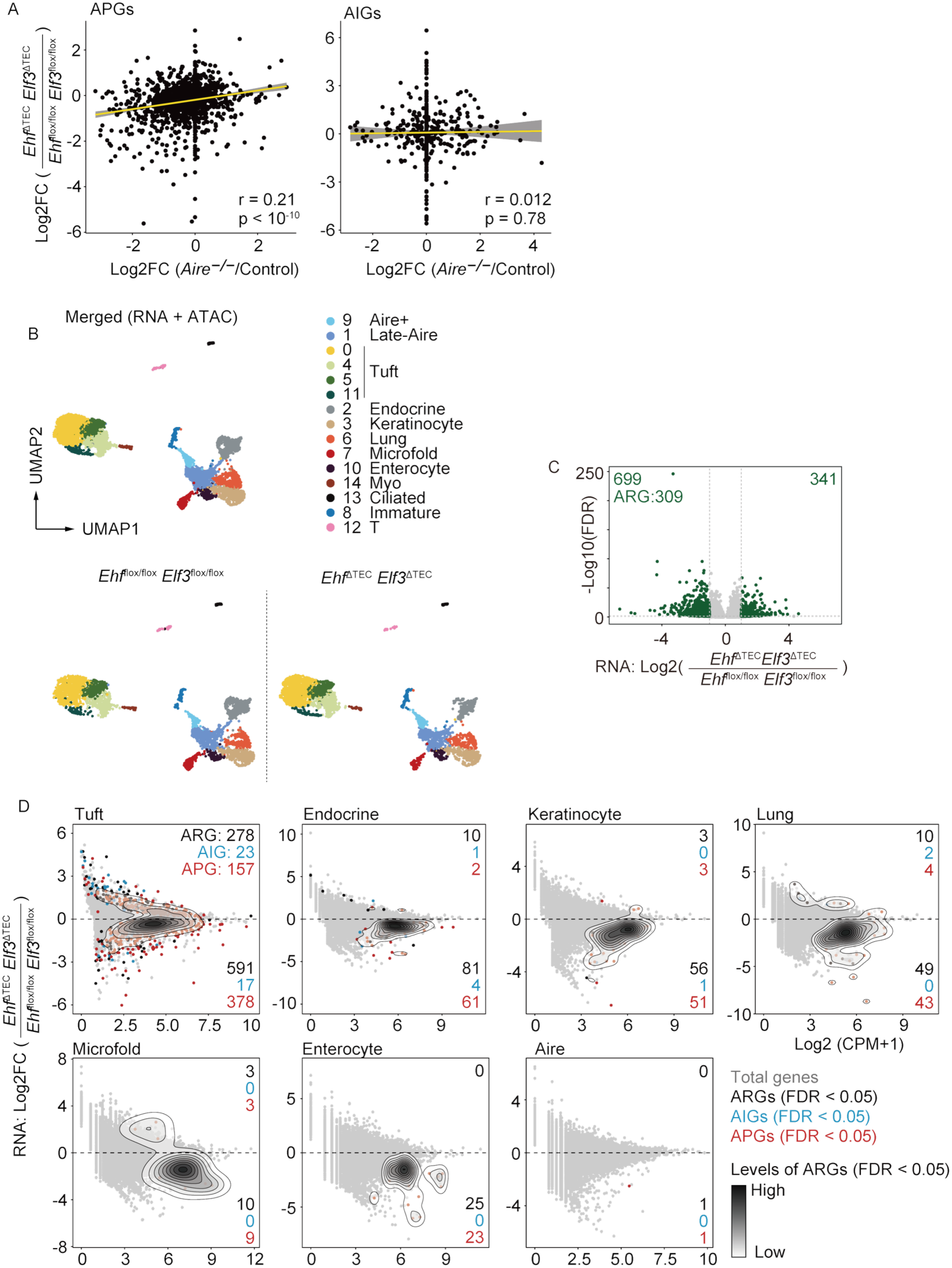
EHF and ELF3 preferentially control the expression of Aire-regulated genes in post-Aire mimetic TECs. A. FC/FC plots of tuft-like post-Aire mimetic TECs comparing *Aire*^−/–^ versus *Ehf* ^ΔTEC^ *Elf3*^ΔTEC^ mice. R- and p-values were calculated using Pearson correlation analysis. B. UMAP for single-nucleus Epi Multiome (RNA + ATAC) after integration of snRNA-seq and scATAC-seq data performed by the weighted-nearest neighbor unsupervised framework. C. Volcano plots for differential expression analysis of whole post-Aire mimetic TECs between *Ehf*^flox/flox^*Elf3*^flox/flox^ and *Ehf*^ΔTEC^*Elf3*^ΔTEC^. Green plots represent significantly changed genes or regions which are defined as FDR < 0.05 and absolute Log2FC > 1. D. MA plots of log-transformed CPM of *Ehf*^flox/flox^*Elf3*^flox/flox^ gene expression and log2 fold changes of gene expression between *Ehf*^flox/flox^*Elf3*^flox/flox^ and *Ehf*^ΔTEC^*Elf3*^ΔTEC^ in indicated cells. Each gray dot represents a gene, and each black dot represents a significantly changed ARGs (FDR < 0.05). The contour reflects the densities of significantly changed ARGs.

Due to the pronounced heterogeneity of post-Aire mimetic TECs and the difficulty of isolating all subtypes by flow cytometry, we next performed single-cell EpiMultiome analysis on TECs enriched for post-AIRE mimetic populations (Supplementary Figure S5) from *Ehf*^ΔTEC^*Elf3*^ΔTEC^ and control *Ehf*^flox/flox^*Elf3*^flox/flox^ mice. Dimensionality reduction and clustering identified 15 TEC clusters, which were annotated based on established marker genes (Figure 4B and Supplementary Figure S5). All clusters were present in both control and *Ehf*^ΔTEC^*Elf3*^ΔTEC^ mice, with minor changes in the relative abundance of certain post-AIRE mimetic TEC subsets (Supplementary Figure S5). Consistent with flow cytometric analysis (Supplementary Figure S3), these results indicate that combined deletion of *Ehf* and *Elf3* does not substantially impair the differentiation or maintenance of post-AIRE mimetic TEC subsets in contrast to the lineage-defining transcription factors^7^.

Differential expression analysis revealed that 699 genes were downregulated in *Ehf*^ΔTEC^*Elf3*^ΔTEC^ post-Aire mimetic TECs, including 309 ARGs (Figure 4C). Examination of ARG expression across individual post-Aire mTEC subsets showed preferential downregulation in multiple populations (Figure 4D), suggesting that EHF and ELF3 contribute to ARG expression across post-Aire mimetic TECs.

Chromatin accessibility analysis revealed that 4,418 regions were less accessible in EHF/ELF3 double-deficient post-Aire mimetic TECs (Figure 5A, B), indicating that these factors help maintain an open chromatin state required for gene expression. Loss of EHF and ELF3 preferentially affected distal regulatory elements rather than TSSs of ARGs (Figure 5C). Among distal elements, those previously bound by AIRE in Aire⁺ mTECs (ABRs) were preferentially affected (Figure 5C), suggesting that these regions are initially primed by AIRE. Consistent with this, EHF- and ELF3-binding motifs were significantly enriched within APG-linked distal regions but not at TSSs (Figure 5D). Moreover, reanalysis of published EHF ChIP-seq data of pan-mTECs^17^ indicated that EHF preferentially binds distal AIRE-bound regions of ARGs compared with distal AIRE-unbound regions (Figure 5E), supporting functional cooperation between AIRE and ETS factors. Interestingly, accessibility at these distal sites was gained in a subset-specific manner among post-Aire mimetic TECs (Figure 5F), indicating cell type–restricted enhancer activation. Together, these findings support a model in which AIRE primes distal regulatory elements of ARGs, enabling subsequent recruitment of EHF and ELF3 to maintain chromatin accessibility and promote transcriptional activation in post-Aire mimetic TECs (Supplementary Figure S6).

**Figure 5.**
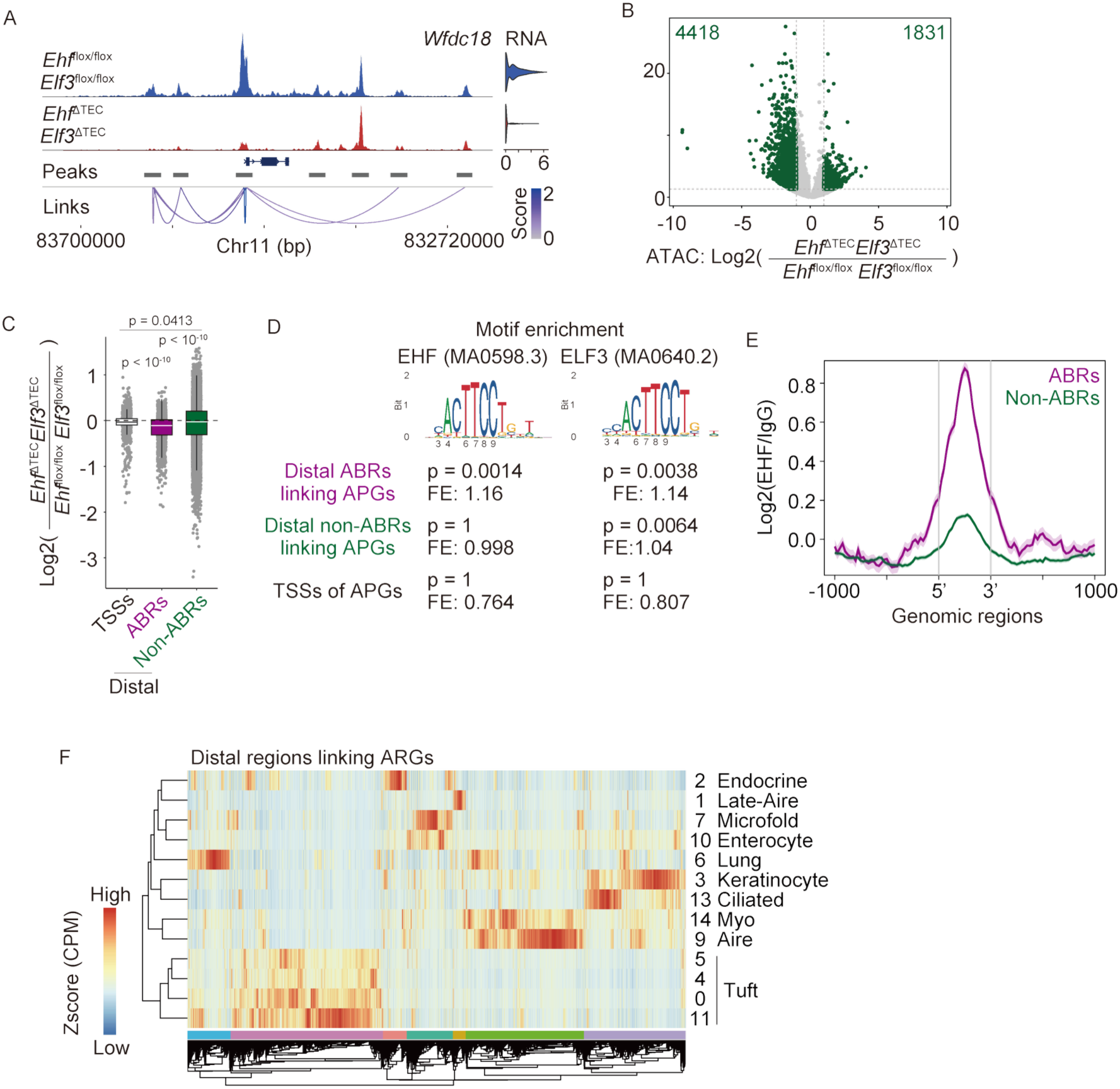
EHF and ELF3 preferentially bind to AIRE-prebound regions and promote their accessibility in post-Aire mimetic TECs. A. Peak tracks of a representative ARG significantly down-regulated by *Ehf*^ΔTEC^*Elf3*^ΔTEC^. Scores mean Z-score calculated by using the correlation coefficient. B. Volcano plots for differential accessibility analysis of whole post-Aire mimetic TECs between *Ehf*^flox/flox^*Elf3*^flox/flox^ and *Ehf*^ΔTEC^*Elf3*^ΔTEC^. Green plots represent significantly changed genes which are defined as FDR < 0.05 and absolute Log2FC > 1. C. Box plots indicating fold changes of APG transcription start sites (TSSs; black) and Aire-bound (purple) or non-Aire-bound (green) distal regions linking APG expression. P-values were calculated by one-way ANOVA with Tukey test. D. The results of motif enrichment analysis for EHF and ELF3 at APG TSSs (black) and Aire-bound (purple) or non-Aire-bound (green) distal regions linking APG expression. E. A profile of mTEC EHF ChIP-seq (Lammers S et al., 2023) at Aire-bound (purple) and non-Aire-bound (green) distal regions linking APG expression. F. A heat map indicating cluster level chromatin accessibility (normalized read counts) at distal regions linking ARG expression.

### Co-expression of EHF and AIRE synergistically enhances ARG expression in cultured cells

To further test whether AIRE priming facilitates EHF/ELF3-mediated gene induction, we transfected HEK293 cells with expression plasmids encoding AIRE-IRES-GFP, EHF-IRES-GFP, both, or a control IRES-GFP vector. GFP⁺ transfected cells were purified by flow cytometry (Supplementary Figure S7) and analyzed by bulk RNA-seq. Overexpression of EHF alone upregulated 30 genes (Figure 6A), including LMO2 and OLFM1, both of which were downregulated in post-AIRE mimetic TECs from *Ehf*^ΔTEC^*Elf3*^ΔTEC^ mice. AIRE expression alone induced five genes (Figure 6B), including KRT5 and CHRNA1, known AIRE targets. Of note, co-expression of AIRE and EHF upregulated 110 genes (Figures 6A, B, Supplementary Figure S7), demonstrating a synergistic effect. Importantly, only the combined expression of AIRE and EHF, but not EHF alone, led to significant enrichment of human orthologs of murine AIRE-dependent genes among the induced set (Figure 6C). These findings support a model in which AIRE primes chromatin to permit EHF-driven gene activation.

**Figure 6.**
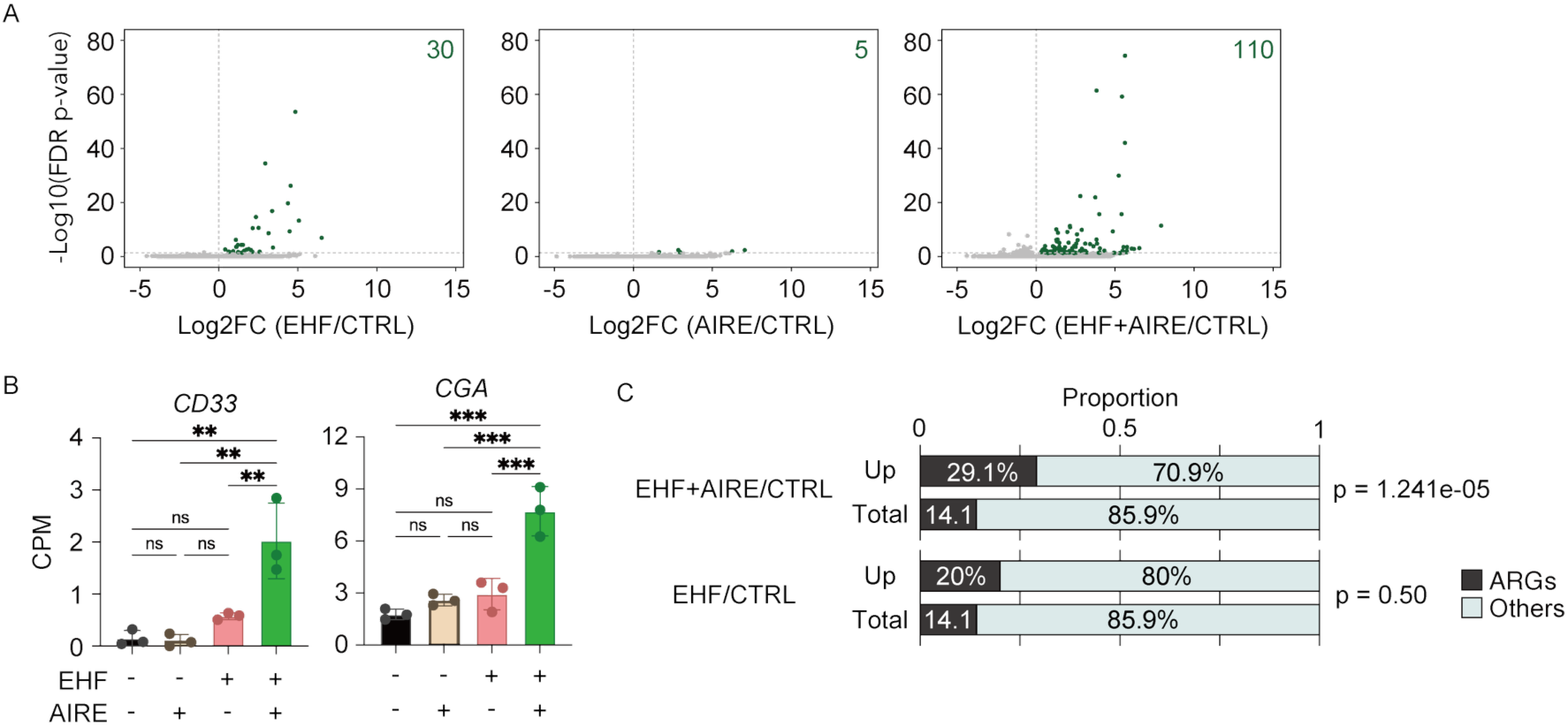
Co-expression of EHF and ELF3 with AIRE increases the expression of a subset of genes in HEK293 cells. A. Volcano plots indicating differentially expressed genes between indicated conditions of over expression in HEK293 cells. Each condition has three biological replicates (N = 3). B. Bar plots indicating expression values of indicated ARG orthologs normalized by 10^6^ read counts in each condition. P-values were calculated using Tukey’s multiple comparisons test. C. Stacking bar plots comparing proportions of ARGs, and non-ARGs in indicated conditions. P-values were calculated using the chai-squared test.

### EHF and ELF3 in TECs preserve self-tolerance in vivo

We next investigated the thymic T cell phenotypes of these mutant mice. Development of conventional thymocytes, Foxp3⁺ regulatory T cells, and their precursors was not significantly affected in *Ehf*^ΔTEC^*Elf3*^ΔTEC^ mice (Supplementary Figure S8). Since post-Aire mimetic TECs promote TSA expression, thereby contributing to thymic self-tolerance, we examined autoimmune manifestations in these mutants. Although splenic activated T cells were not significantly changed, histological analysis of 20-week-old female *Ehf*^ΔTEC^*Elf3*^ΔTEC^ mice revealed marked lymphocytic infiltration in the salivary gland and liver (Figure 7A), along with accumulation of CD3⁺ T cells in the salivary gland (Figure 7B). GSEA analysis suggested that genes associated with salivary gland development were enriched in gene sets reduced in post-Aire mimetic TECs (Figure 7C). However, the commonly targeted tissue in Aire deficiency, the eye, showed no signs of inflammation, nor did the lung (Figure 7A). Overall, these findings suggest a breakdown of self-tolerance causing tissue-selective autoimmunity, partially mirroring the autoimmune phenotypes of *Aire*^−/–^ mice.

**Figure 7.**
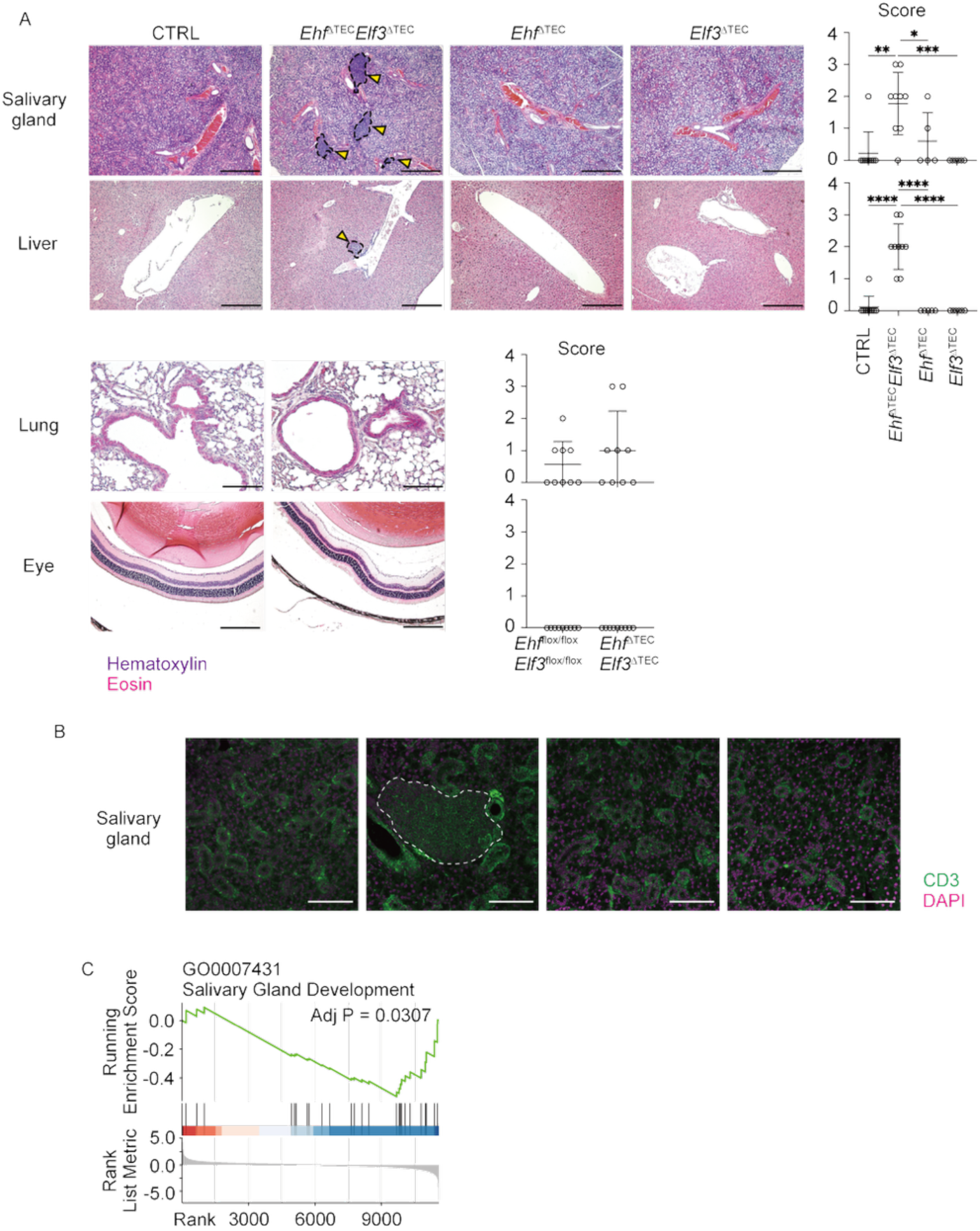
Loss of EHF and ELF3 results in cell infiltration of the salivary gland and liver. A. Representative images for H&E (the region outlined by a dotted line) of salivary gland, liver, lung and eyes in 20-week-old female mice with indicated genotypes. Histological scores reflecting frequences and amounts of lymphocyte infiltration. Bars show mean SEM. P-values were calculated by one-way ANOVA with Tukey test. B. Representative immunohistochemical staining of salivary gland with anti-CD3 antibody. Dot line indicates specific stained area. C. GSEA analysis of the gene ontology term Salivary Gland Development (GO0007431) on differential gene expression profiles of whole post-Aire mimetic TECs between *Ehf^f^*^lox/flox^*Elf3*^flox/flox^ and *Ehf*^ΔTEC^*Elf3*^ΔTEC^ (Figure 4B).

## Discussion

Our study identifies a mechanism by which AIRE activity in Aire⁺ mTECs influences gene expression programs in their post-Aire descendants. While AIRE is well established as a transcriptional regulator that stochastically induces TSA expression in Aire⁺ mTECs^6, 21^, our data suggest that its role is not limited to this transient stage. Specifically, AIRE activity at distal regulatory elements is associated with the subsequent maintenance of ARG expression in post-Aire mimetic TECs. This framework offers an explanation for how ARG expression in post-Aire mimetic TECs can remain dependent on prior AIRE activity despite the loss of AIRE expression.

Our chromatin accessibility analyses suggest that AIRE plays dual roles in regulating the epigenetic landscape. This finding may be consistent with previous findings that AIRE rather close the chromatin for TSA expression in mTECs^28^. On the other hand, scATAC-seq showed that AIRE deficiency leads to reduction of chromatin accessibility in Aire^+^ mTECs. Thus, in addition to promoting accessibility at certain loci, AIRE also appears to preserve other regions in a closed or poised state, which later become accessible during differentiation into post-Aire mimetic TECs. This duality highlights AIRE’s role as a temporal gatekeeper, establishing chromatin states that are selectively interpreted at later stages. Such priming parallels observations in other developmental contexts, where transiently expressed pioneer factors create enduring transcriptional potential^29^, but here is linked directly to the maintenance of immunological tolerance.

Although our data indicate that AIRE primes distal regulatory elements of APGs in Aire⁺ mTECs to enable their subsequent activation in post-Aire mimetic TECs, the precise molecular mechanism underlying this priming remains unclear. AIRE contains several functional domains, including a CARD, SAND, and two PHD fingers. The CARD domain promotes transcription by engaging H3K27ac-marked enhancers and forming condensates at CBP/p300-rich loci^30^, while PHD1 binds unmethylated H3K4^30, 31^, an interaction linked to transcriptional inactivity^30^ but required for subsequent condensate assembly^30^. AIRE also recruits chromatin remodelers^32^, histone modifiers^33, 34^, and transcriptional elongation factors^35, 36^, potentially establishing a permissive chromatin state without fully activating transcription. Alternatively, AIRE may localize to or act at sites of transcription-associated DNA breaks^36, 37^ or non-canonical DNA structures^38^, thereby poising these regions for subsequent engagement by secondary transcription factors. This functional diversity, likely mediated through distinct AIRE domains, suggests a plausible mechanism for AIRE-dependent chromatin priming of APG expression programs. Future work dissecting AIRE’s interaction partners and chromatin-modifying activities at APG loci will be essential to define how AIRE establishes a transcriptionally primed state in Aire⁺ mTECs.

We further identify the ETS family members ELF3 and EHF as critical effectors of this priming mechanism. These transcription factors redundantly sustain the expression of ARGs in post-Aire mimetic TECs, acting on distal regulatory elements primed by AIRE. Our genetic and transcriptional analyses demonstrate that loss of both factors compromises ARG expression and precipitates tissue-selective autoimmunity. Functional synergy between AIRE and EHF/ELF3 observed in heterologous systems reinforces the model of a chromatin relay, in which AIRE primes regulatory elements that are subsequently engaged by ETS factors to secure TSA expression (Supplementary Figure 6).

Variants in AIRE have also been associated with other autoimmune conditions^39^, including rheumatoid arthritis^40^, vitiligo^41^, and systemic lupus erythematosus (SLE) ^42^, indicating that genetic perturbations affecting AIRE function may contribute to autoimmunity beyond classical APECED. Consistent with a broader involvement of thymic transcriptional pathways in autoimmunity, previous GWAS analyses have identified associations between EHF and ELF3 loci and susceptibility to SLE^43, 44, 45^. These population-level correlations do not establish mechanistic causality or clinical overlap with APECED but suggest that variation within this ETS-related transcriptional module may contribute to polygenic autoimmune risk.

Together, our results define a chromatin-based relay mechanism in which AIRE primes distal regulatory elements, and the ETS transcription factors EHF and ELF3 subsequently engage these sites to sustain tissue-specific antigen expression in post-Aire mimetic TECs. This model extends the paradigm of AIRE function beyond transient transcriptional activation to include a lasting priming step that ensures continuity of self-antigen presentation across mTEC differentiation. Disruption of this AIRE–ETS relay results in impaired TSA expression and tissue-selective autoimmunity in mice, and may contribute to autoimmune susceptibility in humans. Future studies will be required to determine how additional transcription factors integrate into this regulatory network and whether similar priming mechanisms operate in the human thymic epithelium.

## Acknowledgment

Computations were partially performed on the NIG supercomputer at ROIS, National Institute of Genetics, and HOKUSAI supercomputer system at RIKEN. This work was supported by cSIMVa Vaccine Challenge Grants.

## Funding

This work was supported by Grants-in-Aid for Scientific Research from JSPS (23K19479 to K.H., 3KK0139 to K.H., 25K18828 to K.H., 21J14033 to K.H., 23K27399 to T.A., 23K06385 to N.A., 24K18386 to T.M), MEXT/JSPS KAKENHI (25H01369 to T.A.), and JSPS KAKENHI Grant Number JP 16H06277(CoBiA) to T.A..

## Author contributions

**Kenta Horie**, Data curation, Formal analysis, Funding acquisition, Investigation, Validation, Writing – original draft and editing; **Yuko Yamagata, Yuki Takakura, Takahisa Miyao, Maki Miyauchi, Tatsuya Ishikawa, Naho Hagiwara, Rin Endo, Hiroto Ishii, Wataru Muramatsu, Fuyoko Suzuki**, Data curation, Validation, Writing – review and editing; **Noritaka Yamaguchi**, Formal analysis, Writing – review and editing; **Nobuaki Yoshida**, Creation of gene-modified mice; **Georg A. Hollander**, Interpretation, Providing materials, Writing – review and editing; **Jun-ichiro Inoue**, Supervision, Providing materials, Writing – review and editing; **Eiryo Kawakami, Shigeo Murata**, Supervision, Writing – review and editing; **Taishin Akiyama, Nobuko Akiyama** Formal analysis, Funding acquisition, Investigation, Project administration, Supervision, Validation, Writing – review and editing

## Animal ethics statement

All animal experiments were conducted at RIKEN in accordance with institutional and national guidelines for the care and use of laboratory animals. Mice were housed in a specific pathogen-free facility at RIKEN with a 12-hour light–dark cycle and ad libitum access to food and water. All procedures were reviewed and approved by the Institutional Animal Care and Use Committee of RIKEN, Yokohama Branch (Protocol No. 2018-075). Experimental procedures were designed to minimize animal suffering, and humane endpoints were employed as necessary. Mice were euthanized using an approved method consistent with the Guidelines for Proper Conduct of Animal Experiments issued by the Science Council of Japan.

## Materials and Methods

### Mice

Female littermates were used for these experiments. Foxn1-Cre mice were reported previously^46^. *Elf3*^flox^ mice on the C57BL/6 background were obtained Jackson Laboratory (B6.Cg-Elf3^tm1Mote^ Tyr^c-2J^/J)^46, 47^. *Ehf*^flox/flox^ mice on the C57BL/6 background were originally generated in our laboratory by using a targeting vector, in which a part of exon 2 of the *Ehf* gene is flanked by LoxP sites. *Elf3*^ΔΤEC^ and/or *Ehf*^ΔΤEC^ mice were generated by crossing Foxn1-Cre mice with *Elf3*^flox^ mice and/or *Ehf*^ΔΤEC^ mice. PCR was performed to identify mouse genotypes by using the following PCR primers that recognize Foxn1^tm3^(cre)^Nrm^ (forward 5’- CAT ACG ATT TAG GTG ACA CTA TAG -3’ and reverse 5’- AAT CTC ATT CCG TTA CGC AG -3’), Elf3^tm1Mote^ (forward 5’- AGT GTT GTG GCC CTG TGT AGT -3’ and reverse 5’-AAC TGA TGG CGA GCT CAG A-3’) and Ehf^flox^ (forward 5’- GGT TGA GGT CCC CAT GAC ATT AC -3’ and reverse 5’- ATC TGA GTA GAT GGC GCT TCA GG -3’).

### Antibodies and reagents for flow cytometry

APC/Cyanine7 anti-mouse CD45 (BioLegend, clone 30-F11, Cat#103116, 1:400), APC/Cyanine7 anti-mouse TER119 (BioLegend, clone TER-119, Cat#116223, 1:200), BV510 anti-mouse EpCAM (CD326) (BioLegend, clone G8.8, Cat#118231, 1:400), FITC anti-mouse EpCAM (CD326) (BioLegend, clone G8.8, Cat#118208, 1:400), PE/Cyanine7 anti-mouse I-A/I-E (BioLegend, clone M5/114.15.2, Cat#107630, 1:3200), Pacific Blue anti-mouse I-A/I-E (BioLegend, clone M5/114.15.2, Cat#107620, 1:3200), Alexa Fluor 647 anti-mouse Ly51 (BioLegend, clone 6C3, Cat#108312, 1:400), PerCP/Cyanine5.5 anti-mouse Ly51 (BioLegend, clone 6C3, Cat#108316, 1:400), PE anti-mouse CD80 (BioLegend, clone 16-10A1, Cat#104708, 1:300), Brilliant Violet 510 anti-mouse CD24 (BioLegend, clone M1/69, Cat#101831, 1:300-400), Brilliant Violet 785 anti-mouse Ly-6A/E (Sca-1) (BioLegend, clone D7, Cat#108139, 1:300), PE anti-mouse CD104 (ITGB4) (BioLegend, clone 346-11A, Cat#123610, 1:400), Brilliant Violet 711 anti-mouse CD104 (ITGB4) (BD, clone 346-11A, Cat#743082, 1:400), FITC anti-mouse Ly-6D (BioLegend, clone 49-H4, Cat#138606, 1:400), PE anti-mouse Podoplanin (BioLegend, clone 8.1.1, Cat#127408, 1:400), Alexa Fluor 647 anti-mouse L1CAM (BioLegend, clone 555, Cat#FAB5674R, 1:50), Biotinylated Ulex Europaeus Agglutinin I (UEA I) (Vector Laboratories, Cat#B-1065, 1:400), PE/Cyanine7 Streptavidin (BioLegend, Cat#405206, 1:800), Alexa Fluor 700 Streptavidin (Invitrogen, Cat# S21383, 1:800), PE/Cyanine7 anti-mouse CD4 (BioLegend, clone RM4-5, Cat#100528, 1:400), PE/Cyanine5 anti-mouse CD4 (BioLegend, clone RM4-5, Cat# 100514, 1:400), APC/Cyanine7 anti-mouse CD8a (BioLegend, clone 53-6.7, Cat#100714, 1:400), Alexa Fluor 594 anti-mouse CD8a (BioLegend, clone 53-6.7, Cat#100758, 1:1000), PE anti-mouse CD25 (BioLegend, clone PC61, Cat#102008, 1:400), FITC anti-mouse CD25 (BioLegend, clone PC61, Cat#102006, 1:400), PE anti-mouse CD44 (BioLegend, clone IM7, Cat#103008, 1:400), Brilliant Violet 421 anti-mouse CD44 (BioLegend, clone IM7, Cat#103040, 1:400), PE anti-mouse TCRγ/δ (BioLegend, clone GL3, Cat#118108, 1:200), APC/Cyanine7 anti-mouse TCR β chain (BioLegend, clone H57-597, Cat#109220, 1:200), FITC anti-mouse CD69 (BioLegend, clone H1.2F3, Cat#104506, 1:400), Alexa Fluor 647 anti-mouse H2-kb (BioLegend, clone AF6-88.5.5.3, Cat#116512, 1:400), APC anti-mouse CD357 (GITR) (BioLegend, clone DTA-1, Cat#126312, 1:400), APC anti-mouse NK-1.1 (BioLegend, clone PK136, Cat#108710, 1:200), FITC anti-mouse/rat/human CD27 (BioLegend, clone LG.3A10, Cat#124208, 1:1000), PE/Cyanine7 anti-mouse CD196 (CCR6) (BioLegend, clone 29-2L17, Cat#129816, 1:50), Alexa Fluor 700 anti-mouse CD103 (BioLegend, clone 2E7, Cat#121442, 1:100), PE anti-mouse FOXP3 (Invitrogen, clone FJK-16s, Cat#12-5773-82, 1:400), Alexa Fluor 647 anti-mouse RORγt (BD, clone Q31-378, Cat#562682, 1:100), PE/Cyanine7 anti-mouse PLZF (BioLegend, clone 9E12, Cat#145806, 1:200), FITC anti-T-bet (BioLegend, clone 4B10, Cat#644812, 1:100), FITC anti-mouse CD62L (BioLegend, clone MEL-14, Cat#104406, 1:400), anti-mouse CD16/32 (BioLegend, clone 93, Cat#101302, 1:200) and PE mCD1d PBS-57 tetramer (NIH)

### Generation of genetically modified mouse at the Ehf locus

*Ehf*^flox/flox^ mice were established by recombination in JM8.A3 ES cells and subsequent injection into blastocysts. A targeting vector at the Ehf locus was commercially obtained from the EUCOMM program (https://www.mousephenotype.org/data/genes/MGI:1270840).

### Cell isolation from murine thymi

Murine thymi from 4-week-old and 8-week-old female mice were machinery dissected using razor blades. Thymic fragments in 1.5 mL tubes were pipetted up and down with 1 ml of HEPES-RPMI (nacalai) and then briefly spun down. Supernatant was collected as lymphocyte fractions to be analyzed. Lymphocyte-removed thymic fragments were dissociated with Liberase (Roche, 0.05 U/mL) plus DNase I (Sigma-Aldrich) in RPMI 1640 by incubation for 12 min at 37 ℃ three times. After removing debris by filtration using 100 µm mesh filters, single cell suspensions were used as stroma cell fraction to be analysed. Live cells and dead cells were counted by AO/PI (Logos Biosystems, Cat#F23001) with LUNA-FL Dual Fluorescence Cell Counter (Logos Biosystems).

### Cell isolation from murine spleens and lymph nodes

Murine spleens and inguinal and brachial lymph nodes from 20-week-old female mice were machinery dissected in 3 mL RPMI 1640 plus 5% FBS using frosted slide glasses. After filtering out debris with 100 µm mesh filters, red blood cells in spleen samples were dissociated by incubation with lysis buffer for 1 min on ice, followed by neutralization by adding 9 mL RPMI 1640 plus 5% FBS. Cells were spun down for 5 min at 1500 ppr and suspended with 1 mL 1xPBS plus 2% FBS. After removing debris by filtration using 100 µm mesh filters, single cell suspensions were used as lymphocytes to be analysed. Live cells and dead cells were counted by AO/PI with the LUNA cell counter.

### Cell preparation for flow cytometric analysis

Cell surface staining was conducted with fluorochrome-labeled antibodies for 20 min on ice in PBS plus 2% FBS following incubation with the CD16/32 antibody to block non-specific bindings of antibodies. Intracellular staining was conducted with kit, according to the manufacture’s instructions. In brief, cells labelled with Zombie Aqua fixable viability kit (BioLegend, Cat#423101, 1:100) in 1x PBS after cell surface staining were fixed permeabilized with permeabilization buffer. These cells were stained with fluorochrome-labelled antibodies for FOXP3, T-bet, PLZF and RORγt in buffer for 20 min on ice. For stroma cell fractions, after cell surface staining, CD45^-^ cells were negatively selected using the magnetic separation technology by MACS with Anti-APC MicroBeads (Miltenyi Biotec, Cat#130-090-855) and LS columns (Miltenyi Biotec, Cat#130-042-401, 1:100). MACS buffer was used to suspend cells and dilute Anti-APC MicroBeads. 7-Aminoactinomycin D (7-AAD) (Nacalai, Cat#19175-34, 1:500) or SYTOX Blue (Invitrogen, Cat#S34857, 1:1000) was supplemented into cell surface-stained samples prior to flow cytometric analysis to gate out 7AAD^+^ or SYTOX Blue^+^ cells as dead cells. Flow cytometry data was analyzed by FlowJo.

### Over expression of Aire and Ehf genes in HEK293 cells

HEK293 cells were cultured in Dulbecco’s modified Eagle’s medium (DMEM) supplemented with 10% FBS, L-glutamate and penicillin/streptomycin antibiotics at 37 °C with 5% CO2. Aire and Ehf were expressed in HEK293 cells by transfection of different plasmids, respectively. Plasmids driving expression of mouse Aire or Ehf were constructed by in-frame insertion at an EcoRI cite of the pMX-IRES-GFP, according to the manufacturer’s instructions (TAKARA, In-Fusion® HD Cloning Kit, Cat#639648). Indicated plasmids were transfected into cells on 6-well plates using Lipofectamine3000.

### Western blotting for HEK293 cells expressing AIRE and EHF

GFP^+^ cells were collected into RPMI plus 5% FBS by FACS AriaIII (BD) with the yield option as Aire and/or Ehf expressing cells. Cells lysed with Laemmli buffer were subjected to SDS-polyacrylamide gel electrophoresis and electrotransferred onto polyvinylidene difluoride membranes. Transferred membranes were incubated with antibodies (EHF, AIRE, ACTIN) at 4C overnight to detect indicated proteins which were captured using a ChemiDoc XRS+ image analyzer (Bio-Rad). Band intensity was measured by Quantity One software (Bio-Rad). The following antibodies were used for western blot: EHF polyclonal antibody (Abcam, Cat#Ab105375, 1:1000), anti-mouse AIRE (Thermo Fisher Scientific, clone 5H12, 14-5934-82, 1:500), anti-mouse β-Actin (MBL, clone 6D1, M177-3, 1:1000)

### Library preparation for bulk RNA-seq

Bulk RNA-seq for murine samples was conducted regarding sub populations of TECs (CD45^-^TER119^-^ EpCAM^+^ cells). Immature mTEC (UEA-1^+^ MHCII^lo^ Ly51^-^ ITGB4^+^ TECs), Aire^+^ mTEC (UEA-1^+^ MHCII^hi^ Ly6D^lo^ TECs), post-Aire mTEC (UEA-1^+^ CD80^hi^ CD24^+^ ScaI^+^ TECs), Tuft-like mTEC (UEA-1^+^ MHCII^lo^ Ly51^-^ L1CAM^+^ TECs) and cTEC (UEA-1^-^ Ly51^+^ MHCII^hi^ TECs) were sorted directly into cell lysis buffer containing 2xTCL and 2-ME by FACS AriaIII (BD). After sorting cells, the concentration of TCL was adjusted to 1x TCL with nuclease-free water or cell lysis buffer depending on how many volumes were obtained during cell sorting. RamDA-seq technology was applied for bulk RNA-seq analysis to profile transcriptome from small number of cells. Libraries were generated according to the protocols produced by the original paper of RamDA-seq^48^. In brief, after purification of nucleic acids with Agencourt RNA Clean XP (Beckman Coulter) and subsequent treatment with DNase I, the RT-RamDA mixture containing 2.5× PrimeScript Buffer (TAKARA), 0.6 µM oligo(dT)18 (Thermo Fisher Scientific), 10 µM 1st NSR primer mix, 100 µg/mL of T4 gene 32 protein, and 3× PrimeScript enzyme mix (TAKARA) were added to the purified nucleic acids for reverse transcription. Samples were added to second-strand synthesis mix containing 2× NEB buffer 2 (NEB), 625 nM dNTP Mixture (NEB), 25 µM 2nd NSR primers, and 375 U/mL of Klenow Fragment (3’–5’ exo-) (NEB). After cDNA synthesis and subsequent purification by AMPure XP (Beckman Coulter), sequencing library DNA was prepared using the Tn5 tagmentation-based method. Bulk RNA-seq was also conducted regarding HEK293 cells over expressing Aire and/or Ehf genes. GFP^+^ HEK293 cells were sorted into RPMI plus 5% FBS by FACS AriaIII (BD). RNA was extracted with Trizol (Invitrogen). Construction of RNA library and sequencing was conducted by Novogene Co., LTD (Beijing, China). Sequencing was performed by a DNBSEQ T7 (MGI).

### Data analysis for bulk RNA-seq

For libraries from murine samples, FASTQ files by paired-end sequencing on HiSeqX (Illumina) were processed with FASTP program to trim adaptor sequences^49^. Read alignment with the mouse genome reference mm10, read count calculation, read count normalization and differential gene expression analysis were performed by CLC main bench workstation (Version 7.5.1; Qiagen). Genes with a mean log-normalized expression value below 1 across all samples were excluded from downstream analysis. Gene set enrichment analysis (GSEA) was performed using the fgsea package (version 1.30) in R.

For libraries from the HEK293 cell line, FASTQ files by next generation sequencing were processed with FASTQC (version 0.12.1) to check qualities of libraries and FASTP (version 1.0.1) to trim adaptor sequences. Read alignment with the human reference HG39 was performed with STAR (version 2.7.11b)^50^. Aligned reads with under 30 MAPQ score, and duplicate reads were removed by the SAMtools^51^ and PICAD (https://broadinstitute.github.io/picard/; version 3.2.0) program, respectively. Passed reads were counted every gene with HTSeq-count (version 2.0.5)^52^ and read count tables of each sample were combined and processed with DESeq2^53^ (version 1.44) in R (version 4.4.1) language to detect the changes in gene expression.

### Data processing for single-cell RNA-seq of^-^ TECs

FASTQ files from Aire^+/+^(control) TEC libraries and Aire^−/–^ TEC libraries publicly obtained from the Gene Expression Omnibus (GEO) under accession code GSE206886 were aligned to the mm10 references with Cell Ranger. Output files from Cell Ranger were processed to remove low-quality cells, normalize read counts and integrate data for further downstream analyses with the Seurat (version 5.3)^54^ in R (version 4.4.1). Low-quality cells were defined based on UMI counts and percentages of mitochondrial RNA. Data integration among samples was conducted using the RPCA integration method based on principal component analysis (PCA) which had been conducted with the 3000 most variable genes selected with variance stabilizing transformation prior to data integration. After integration of four independent data sets, Uniform Manifold Approximation and Projection (UMAP) followed by shared-nearest-neighbor graph construction was performed with the top 30 PCs. The resolution of cell clustering analysis was set at 0.5. Subclustering analysis was conducted, which were associated with Aire^+^, late-Aire, and post-Aire mimetic TECs (Cluster 0,2,5,6,7,9,11,13,16,18 in Supplementary Figure. 1). The integrated dataset was split into four samples at once, and these samples were then re-integrated using the same strategy as the integration of total TECs described above.

### Data analysis for single-cell RNA-seq of TECs

Monocle2 (version 2.32.0) was applied to the integrated dataset of these mTECs to construct tree graphs by the DDRTree method and predict pseudotime, according to the instructions provided by Monocle2 developers^25^. Branched expression analysis modeling (BEAM)^55^ was performed to analyze the branching points and detect differentially expressed genes between the branches. Differential expressed gene analysis between genotypes along the pseudotime trajectory was performed with the differentialGeneTest function.

### Data processing and analysis for single-cell ATAC-seq of TECs

FASTQ files from two control and two *Aire*^−/–^ TEC libraries publicly obtained from the Gene Expression Omnibus (GEO) under accession code GSE206886 were aligned to the mm10 references and peaks were then detected by Cell Ranger ATAC (version 2.1.0). FASTQ files of scEpiMultiome for wild-type TEC libraries publicly obtained under GSE236667 were aligned to the mm10 references and peaks were then detected by Cell Ranger ARC (version 2.0.2). Peak files from each sample were combined into one peak file to be used for integrative analysis. Seurat objects created from Cell Ranger output files based on a combined peak file were processed to remove low-quality cells, normalize read counts through Term Frequency-Inverse Document Frequency (TF-IDF) transformation and integrate data for further downstream analyses with the Seurat (version 5.3) and Signac (version 1.14)^54^ in R. Low-quality cells were defined based on filter-passed read counts and percentages of filter-passed reads mapped in detected peak regions to total filter-passed reads. Prior to data integration, singular value decomposition was performed to reduce dimensions in latent semantic indexing (LSI). Data integration among samples was conducted based on anchors which was selected with robust LSI on dimensions 2 through 50. After integration of four independent data sets, UMAP followed by shared-nearest-neighbor graph construction was performed incorporating dimensions 2 through 50. The resolution of cell clustering analysis was set at 0.8. Subclustering analysis was associated with Aire^+^, late-Aire, and post-Aire mimetic TECs (Cluster 1,4,5,6,9,11,13,14,15,16,17,18). Differentially accessible regions between indicated populations were detected by Wilcox rank test and BH p-value adjustment method. Enriched motifs in indicated regions compared to background peaks were calculated by ChromVAR^56^ (version 1.26) function with the JASPER 2022 motif database. Peaks correlated with gene expression were identified using the LinkPeaks function implemented in the Signac package. Promoters were defined as regions within ±500 bp of transcription start sites (TSSs).

### Data analysis for public ChIP-seq datasets

Datasets of ChIP-seq for EHF and AIRE were publicly obtained from the Gene Expression Omnibus (GEO) under accession code (GSE92597 for AIRE ChIP-seq from mTECs, GSE232702 for EHF ChIP-Seq from mTECs). FASTQ files of these were processed with FASTQC to check quality and FASTP to trim adaptor sequences prior to alignment. BOWTIE2 (version 2.5.4)^57^ was used for read alignment to the mm10 reference. After removal of low-quality reads with MAPQ score under 30, peak calling was performed by MACS3 (version 3.0.1)^58^. AIRE-binding regions were identified by using the irreproducible discovery rate (IDR) framework (version 2.0.4.2). Peaks with an IDR score > 540 were defined as reproducibly detected peaks between two AIRE ChIP-Seq datasets. NGS plot was used to show distribution of mapped reads in indicated genomic regions^59^.

### Library preparation for droplet-based single-nuclear EpiMultiome

Mimetic cells (MHCII^lo^ Ly51^-^ ITGB4^-^ Podoplanin^-^ TECs) from 4-week-old *Ehf^f^*^lox/flox^*Elf3*^flox/flox^ or *Ehf*^ΔTEC^*Elf3*^ΔTEC^female mice were sorted into RPMI 1640 plus 5% FBS by FACS AriaIII (BD). Three thymi were pooled every genotype and used as one sample after cell sorting to obtain enough cells. Sorted cells were spun down for 5 min at 300 xg and suspended with 2% after discarding supernatant. After counting cell number, cells were incubated with 100 µl cell lysis buffer for 3 min on ice to extract intact nuclei after discarding supernatant following centrifuge for 5 min at 300 xg. After incubation, 1mL buffer was immediately added and pipetted up and down 10 times to neutralize cell lysis buffer. After 2 times of wash with buffer, nuclei quality was checked under microscope and nuclei concentration was adjusted to nuclei/µl with nuclei buffer containing. Adjusted nuclei were processed with 10x ChromiumX to generate emulsion in which each droplet included one nuclear and one cell barcoding bead. Libraries were generated using Chromium Next GEM Single Cell Multiome ATAC + Gene Expression Reagent Kit Chemistry, according to the manufacturer’s instructions.

### Data processing and analysis for single-nuclear EpiMultiome

FASTQ files from *Ehf*^flox/flox^*Elf3*^flox/flox^ mimetic and *Ehf*^ΔTEC^*Elf3*^ΔTEC^mimetic TEC libraries by paired-end sequencing on an Illumina were aligned to the mm10 references and peaks were then detected by Cell Ranger ARC. Peak files from each sample were combined into one peak file to be used for integrative analysis. Seurat objects containing RNA counts and ATAC counts created from Cell Ranger ARC output files given a combined peak file were processed to remove low-quality cells, normalize read counts and integrate data for further downstream analyses with the Seurat V5 and Signac package^54^ in R. Low-quality cells were filtered out to retain cells with 10^3^-10^5^ filter-passed ΑTAC read counts, 50-90% of filter-passed ATAC reads mapped to detected peak regions in total filter-passed ATAC reads, a minimum of 1500 UMI counts and a maximum of 20% on mitochondrial RNA. Integration of snRNA-seq datasets was performed using the same strategy with top 50 PCs as described above in ‘Data analysis for single-cell RNA-seq of control and Aire^−/–^ TECs’. Integration of snATAC-seq datasets was also performed using the same strategy as described above. snRNA-seq and snATAC-seq data were integrated and comprehensively analyzed using the Weighted Nearest Neighbor (WNN) method. The resolution of cell clustering analysis was set at 0.7. Differentially accessible regions between indicated populations were detected by Wilcox rank test and BH p-value adjustment method. Enriched motifs in indicated regions compared to background peaks were calculated by ChromVAR function with the JASPER 2022 motif database. Peaks correlated with gene expression were identified using the LinkPeaks function implemented in the Signac package. Promoters were defined as regions within ±500 bp of transcription start sites (TSSs). Gene set enrichment analysis (GSEA) was performed using the fgsea package (version 1.30) in R.

### Histology analysis for autoimmunity monitorin***g***

Tissues from 20-week-old Ehf^flox/flox^ Elf3^flox/flox^ and Ehf^ΔTEC^ Elf3^ΔTEC^ female mice were fixed in 4% formaldehyde (MildformR 10N, Wako, Cat#133-10311) for T cell infiltration analysis. 5 µm sections from paraffin-embedded tissues were stained with eosin (Wako) and hematoxylin (Wako) according to standard protocols. Immunofluorescence staining was performed with CD3 polyclonal antibody (Proteintech, Cat#17617-1-AP). In brief, deparaffinized 5 µm sections from FFPE tissues were treated with a retrieval solution (BD, Retrievagen A, Cat#550524) for 10 min at 90 °C. Tissue sections were incubated with diluted anti-CD3 antibody (1:100) for 2 hours at RT followed by incubation with Alexa Fluor 488 Streptavidin (Thermo Fisher Scientific, Cat#S32354, 1:300) and 50 µg/ml DAPI (vector lab) for 40 min at RT. Images were acquired on a SP8 LIGHTNING (Leica). In general, scores from 0 to 4 indicate severity of lymphocytic infiltration, and complete destruction, respectively, according to the previous report^60^. 0: no detectable infiltration; 1: a focus of perivascular infiltration; 2: several foci of perivascular infiltration; 3: cellular infiltration in less than 50% of the vasculature; 4: cellular infiltration in more than 50% of the vasculature.

### Statistics

Statistical analyses were performed using R (version 4.4.1), CLC Genomics Workbench (Qiagen) and GraphPad Prism10 in this study. We applied the negative binomial generalized linear model, Wilcoxon rank-sum test, logistic regression framework, hypergeometric test, chi-squared test, or one-way ANOVA followed by Tukey’s multiple comparison test. P-values were corrected for multiple testing using the Benjamini-Hochberg or Bonferroni method. The specific statistical methods used for each analysis are detailed in the corresponding figure legends.

### Data and code availability

Sequencing data were deposited in GSE316885 for 10x single-cell Epi Multiome data (scEpiMultiome), GSE316886 for bulk RNA-seq of ETS TECs, and GSE316883 for bulk RNA-seq of HEK293. Public data sets reanalyzed in this paper were obtained from GEO via the following accession codes: GSE92597 for AIRE ChIP-seq from mTECs, GSE232702 for EHF ChIP-seq from mTECs, GSE236667 for scEpiMultiome of wilt-type mTECs and GSE206886 for scRNA-seq, scATAC-seq and bulk RNA-seq from control and Aire^−/–^ TECs. The codes and scripts used in this study are available at GitHub: https://github.com/tken18/ETS_MimeticCells

## Notes

### Competing Interest Statement

The authors have declared no competing interest.

