## Supplementary Figure for "A chromatin relay from AIRE to ETS transcription factors sustains peripheral antigen expression in the thymic mimetic cells to ensure central tolerance"

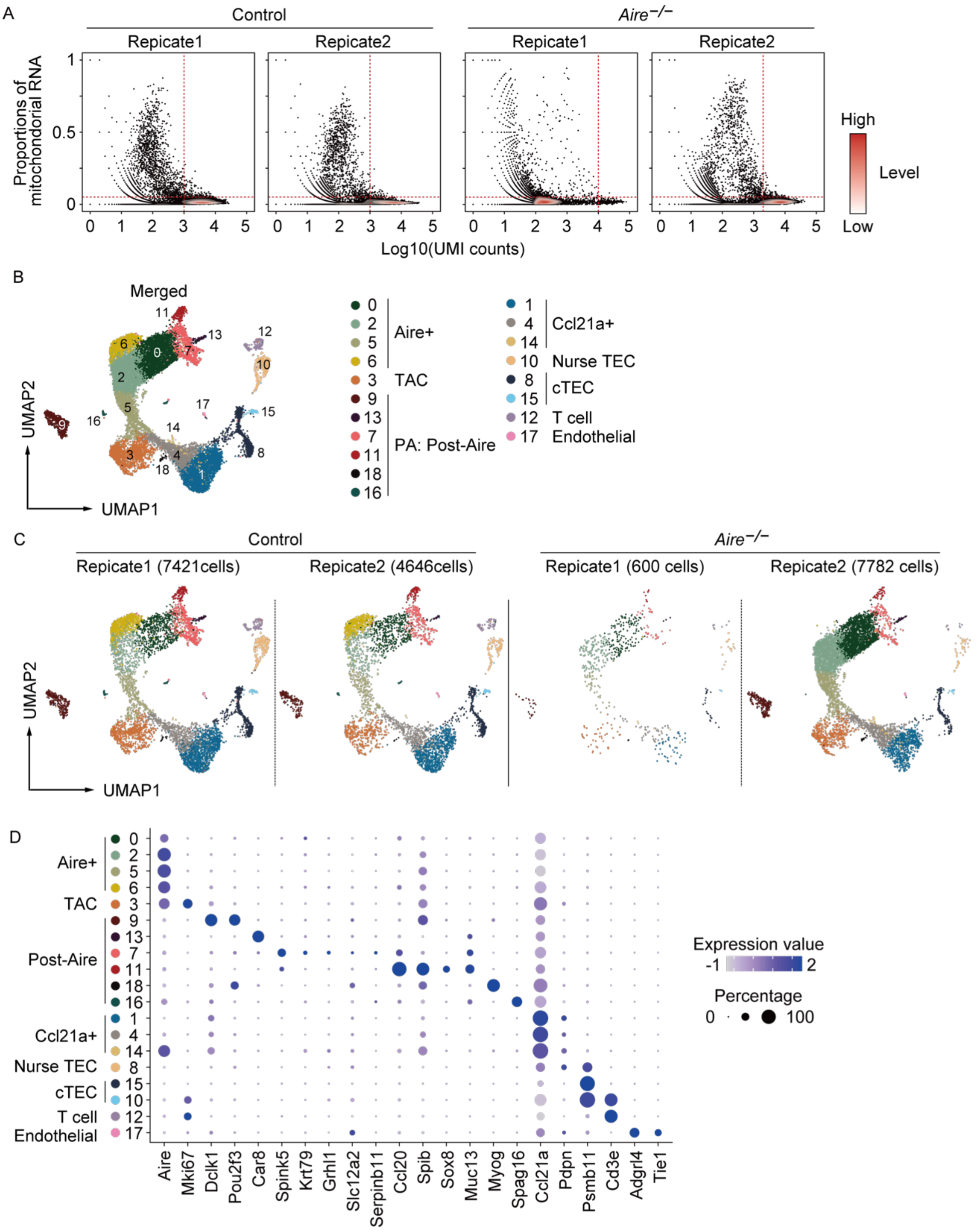


**Supplementary Figure S1. scRNA-Seq analysis on whole TECs from Control and *Aire*$^{-/-}$**

A. Scatter plots indicating proportions of mitochondrial RNA and UMI counts for each scRNA-Seq library. Each dot represents one cell, and red lines indicate threshold for filtering low-quality cells.
B. Merged UMAP of scRNA-Seq on all TECs.
C. UMAP split by samples.

D. Heat dot plot of marker genes for each cell types.

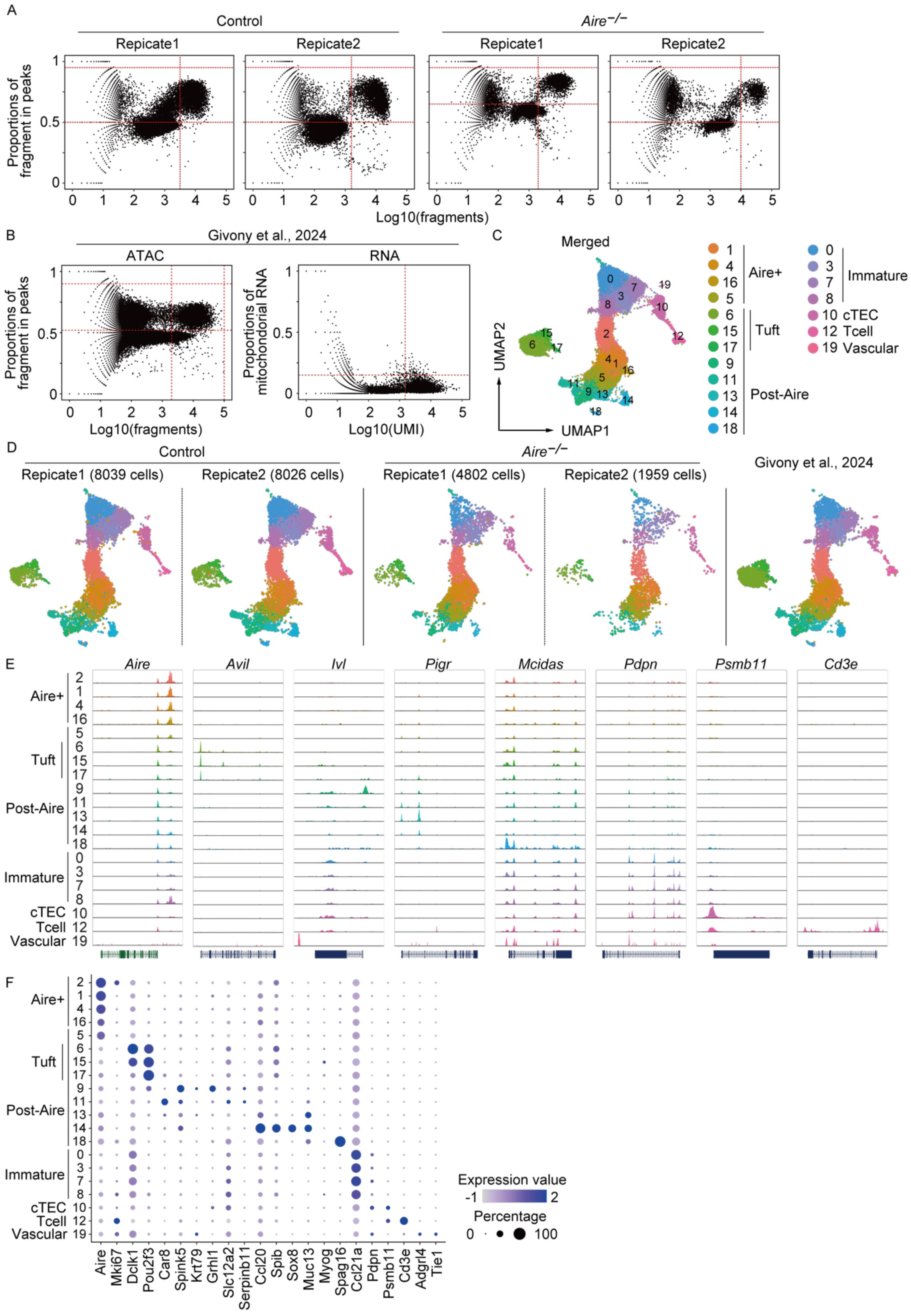


**Supplementary Figure S2. Integration analysis of scATAC-Seq on whole TECs from**

**Control and *Aire*$^{-/-}$ with TEC scEpiMultiome data from Givony et al., 2024.**

A. Scatter plots indicating proportions of fragment in peaks and fragments for each scATAC-Seq library. Each dot represents one cell, and red lines indicate threshold for filtering low-quality cells.
B. Scatter plots for scEpiMultiome data (Givony et al., 2024). A left scatter plot indicating proportions of fragment in peaks and fragments for each scATAC-Seq library. Each dot represents one cell, and red lines indicate threshold for filtering low-quality cells. A right scatter plot indicating proportions of mitochondrial RNA and UMI counts for each scRNA-Seq library. Each dot represents one cell, and red lines indicate threshold for filtering low-quality cells.
C. Merged UMAP of all samples.
D. UMAP split by samples.
E. Peak tracks of genome regions at marker gene for each cell types.
F. Heat dot plot of marker genes for each cell types from scEpiMultiome data (Givony et al., 2024).

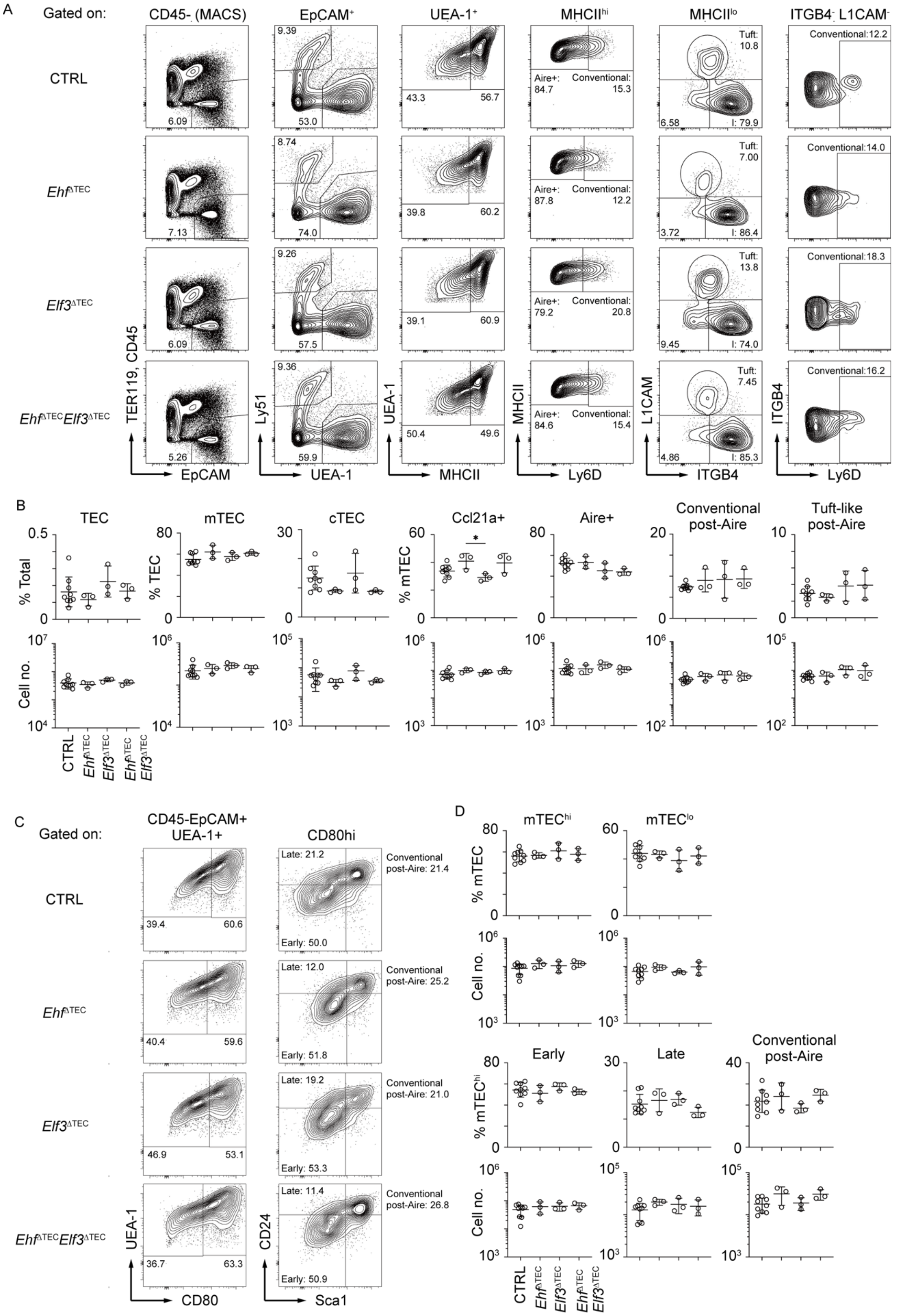


**Supplementary Figure S3. FACS analysis on TEC subsets**

A,C. Gating strategy for TEC subpopulations in the thymic cells from control, *Ehf*$^{\Delta TEC}$, *Elf3*$^{\Delta TEC}$, and *Ehf*$^{\Delta TEC}$*Elf3*$^{\Delta TEC}$ 4-week-old mice. $CD45^{-}$ and $TER119^{-}$ cells are enriched by MACS prior to FACS analysis.

B,D. Cell number and percentages of indicated TEC subpopulations in control (N = 9), *Ehf*$^{\Delta TEC}$ (N = 3), *Elf3*$^{\Delta TEC}$ (N = 3) and *Ehf*$^{\Delta TEC}$*Elf3*$^{\Delta TEC}$ mice (N = 3). Each circle indicates one mouse, and bars show mean ± SEM. *P < 0.05; Turkey-Kramer test.

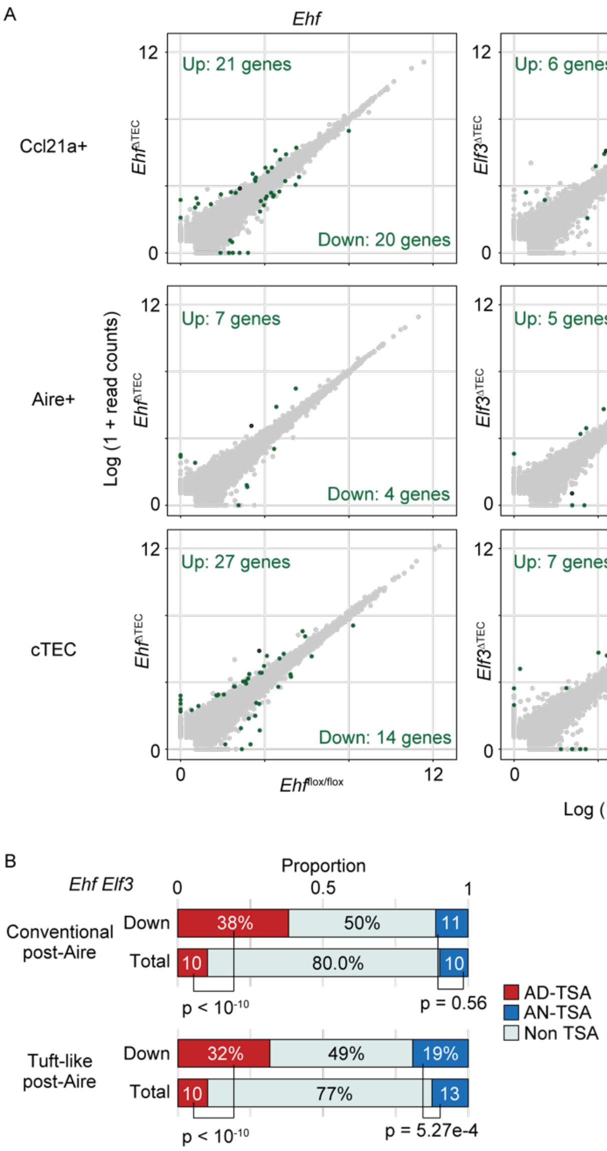


**Supplementary Figure S4. Bulk RNA-Seq analysis on TEC subsets**

A. Scatter plots comparing log-normalized gene expression values of TEC subpopulations in indicated genotypes. Each dot represents a gene, and green plots indicates significantly altered genes, defined by absolute fold change > 2 and FDR < 0.05.

B. Bar plots comparing proportions of AIRE-dependent TSA genes, AIRE-independent TSA genes, and non-TSA genes in indicated mTEC subpopulations. TSA types were defined based on the previous study (Sansom et al., 2014). P-values were calculated using the chai-squared test.

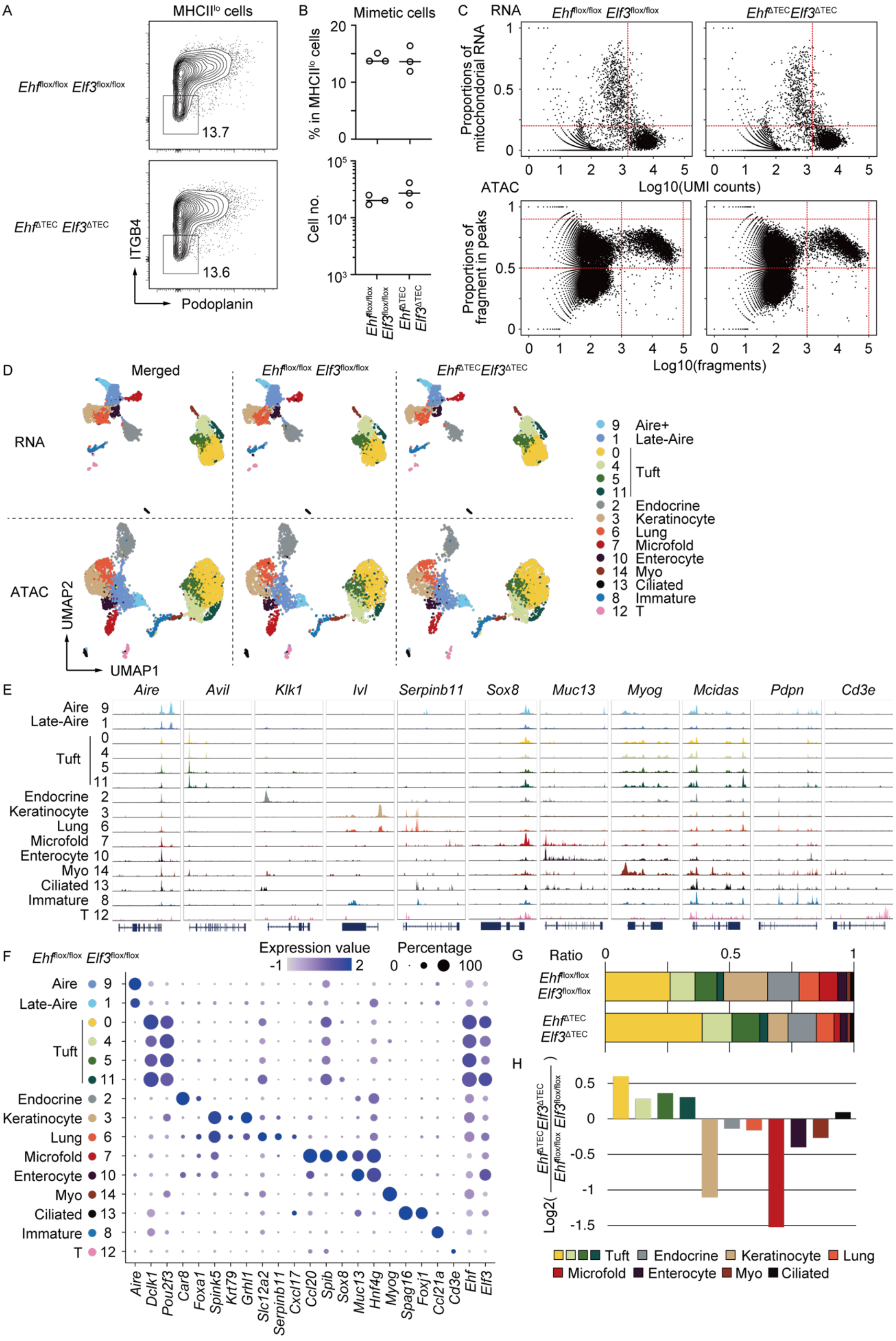
A
MHCIIlo cells
Ehfflox/flox Elf3flox/flox
EhfΔTEC Elf3ΔTEC
13.7
13.6
ITGB4
Podoplanin
B
Mimetic cells
% in MHCIIlo cells
Cell no.
Ehfflox/flox Elf3flox/flox
EhfΔTEC Elf3ΔTEC
C
RNA
Proportions of mitochondorial RNA
Log10(UMI counts)
ATAC
Proportions of fragment in peaks
Log10(fragments)
D
Merged
RNA
ATAC
UMAP1
UMAP2
9 Aire+
1 Late-Aire
0 4 5 11 Tuft
2 Endocrine
3 Keratinocyte
6 Lung
7 Microfold
10 Enterocyte
14 Myo
13 Ciliated
8 Immature
12 T
E
Aire Avil Klk1 Ivl Serpinb11 Sox8 Muc13 Myog Mcidas Pdpn Cd3e
F
Expression value
Percentage
Aire Dclk1 Pou2f3 Car8 Foxa1 Spink5 Krt79 Grhl1 Slc12a2 Serpinb11 Cxcl17 Ccl20 Spib Sox8 Muc13 Hnf4g Myog Spag16 Foxj1 Ccl21a Cd3e Ehf Elf3
G
Ratio
H
Log2(EhfΔTEC Elf3ΔTEC / Ehfflox/flox Elf3flox/flox)
Tuft
Endocrine
Keratinocyte
Lung
Microfold
Enterocyte
Myo
Ciliated

**Supplementary Figure S5. scEpiMultiome analysis of post-Aire mimetic TECs from *Ehf*$^{flox/flox}$ *Elf3*$^{flox/flox}$ and *Ehf*$^{\Delta TEC}$ *Elf3*$^{\Delta TEC}$**

A. Gating strategy for mimetic TECs in the thymic cells from *Ehf*$^{flox/flox}$*Elf3*$^{flox/flox}$ and *Ehf*$^{\Delta TEC}$*Elf3*$^{\Delta TEC}$ mice.
B. Cell number and percentages of mimetic TECs in *Ehf*$^{flox/flox}$*Elf3*$^{flox/flox}$ (N = 3) and *Ehf*$^{\Delta TEC}$ *Elf3*$^{\Delta TEC}$ mice (N = 3). Each circle indicates one mouse, and bars show means.
C. Upper scatter plots indicating proportions of mitochondrial RNA and UMI counts for each snRNA-Seq library. Lower scatter plots indicating proportions of fragment in peaks and fragments for each scATAC-Seq library. Each dot represents one cell, and red lines indicate threshold for filtering low-quality cells. Each dot represents one cell, and red lines indicate threshold for filtering low-quality cells.
D. Merged and split UMAP of snRNA-Seq and scATAC-Seq from mimetic cells from *Ehf*$^{flox/flox}$*Elf3*$^{flox/flox}$ and *Ehf*$^{\Delta TEC}$*Elf3*$^{\Delta TEC}$ mice.
E. Peak tracks of genome regions at marker gene for each cell types.
F. A heat dot plot of marker genes for each cell types from snRNA-Seq data of *Ehf*$^{flox/flox}$*Elf3*$^{flox/flox}$ and *Ehf*$^{\Delta TEC}$*Elf3*$^{\Delta TEC}$ mice.
G. Bar plots of cell proportions in indicated genotypes.
H. Bar plots showing log2 fold changes of cell proportions between *Ehf*$^{flox/flox}$*Elf3*$^{flox/flox}$ and *Ehf*$^{\Delta TEC}$*Elf3*$^{\Delta TEC}$ about indicated mimetic cell populations.

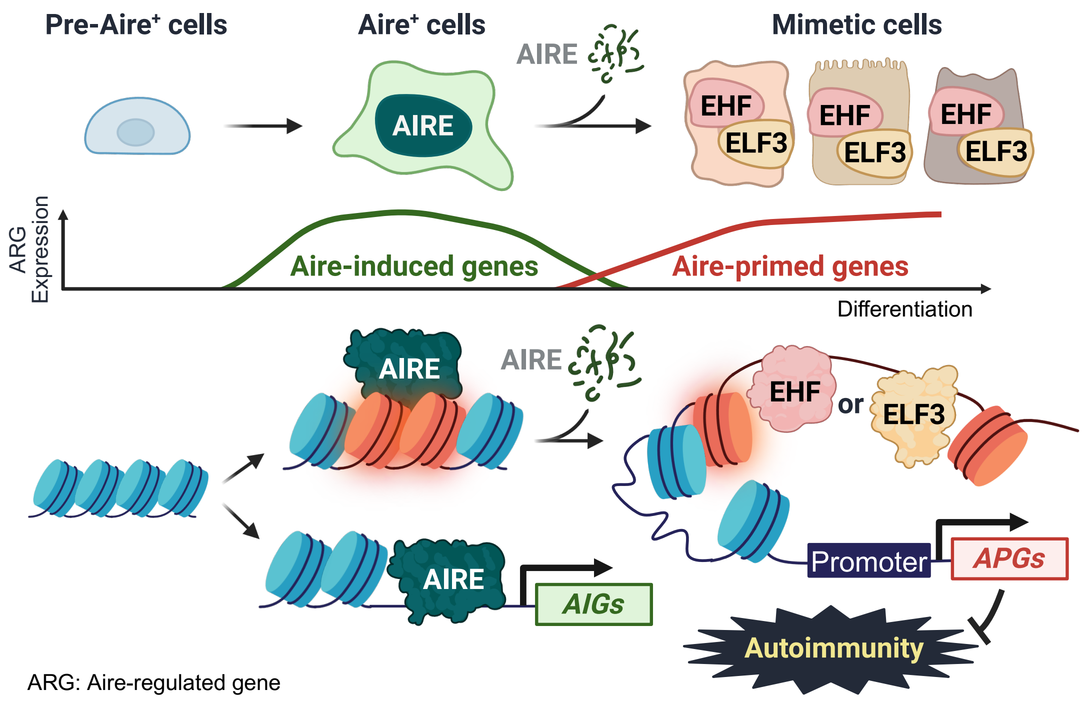


**Supplementary Figure S6. Proposed mechanism of ETS-dependent ARG expression in post-Aire mimetic TECs**

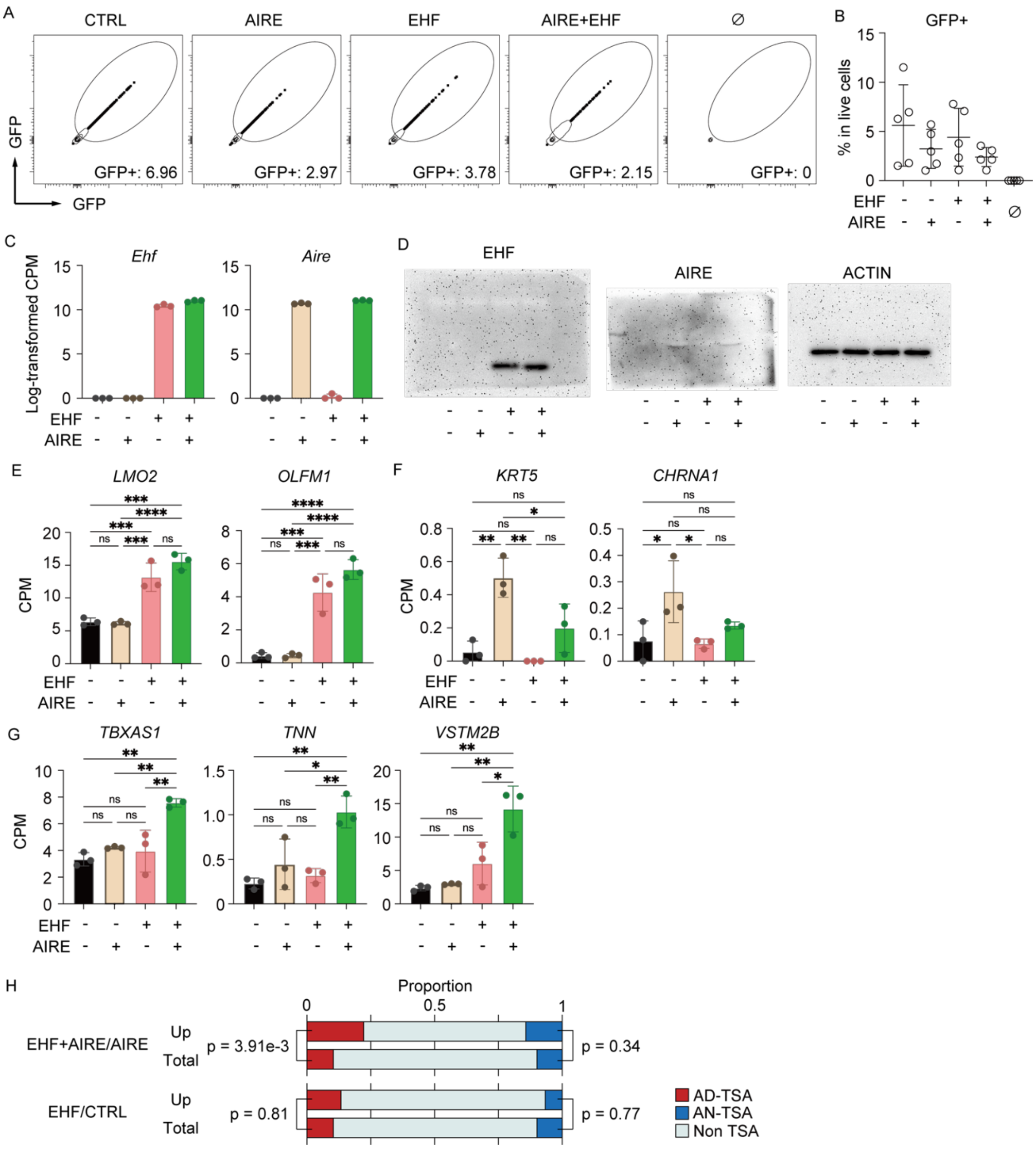


**Supplementary Figure S7. Bulk RNA-Seq on HEK293 cells expressing Aire and Ehf**

A. Gating strategy of $GFP^+$ HEK293 cells.

B. Proportions of $GFP^+$ cells in indicated cell types. No significant change was found by Tukey's multiple comparisons test.

C. Bar plots indicating expression values of mouse Aire and Ehf genes normalized by $10^6$ read counts in each condition. The mouse mm10 reference was used to calculate read counts.

D. Western blotting of indicated mouse proteins for each cell type.

E-G. Bar plots indicating expression values of indicated genes normalized by $10^6$ read counts in each condition. P-values were calculated using Tukey's multiple comparisons test.

H. Stacking bar plots comparing proportions of AIRE-dependent TSA genes, AIRE-neutral TSA genes, and non-TSA genes in indicated conditions. TSA types were defined based on the previous study (Sansom et al., 2014). P-values were calculated using the chai-squared test.

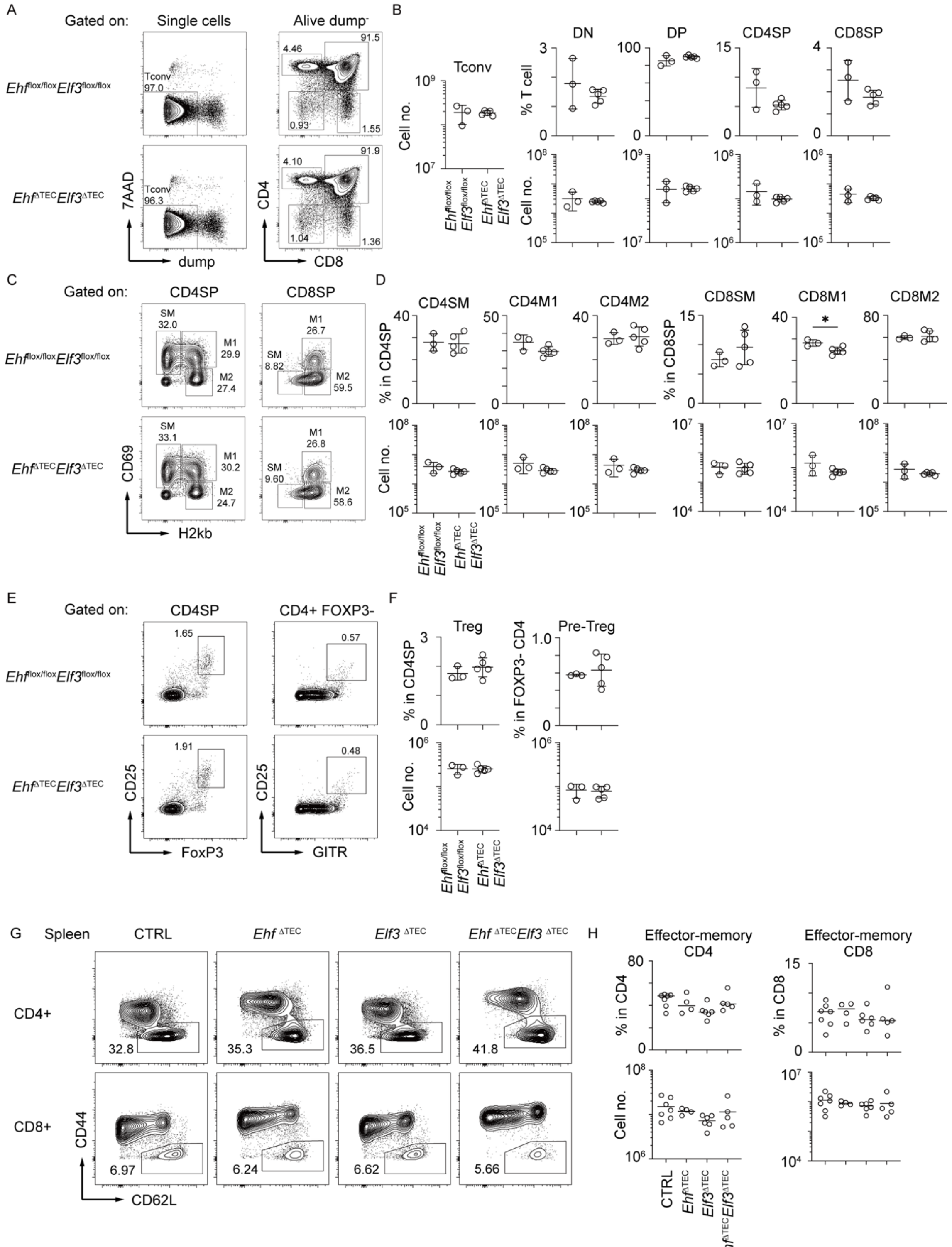


**Supplementary Figure S8. FACS analysis on T cells in the thymus and spleen**

A-F. FACS analysis on thymocytes from 8-week-old *Ehf*$^{flox/flox}$ *Elf3*$^{flox/flox}$ (N = 3) and *Ehf*$^{\Delta TEC}$ *Elf3*$^{\Delta TEC}$ (N = 3) female mice. Each circle indicates one mouse, and bars show mean ± SEM. No significant difference was found by T-test.

A. Representative FACS profiles for CD4 and CD8 expression from 8-week-old mice. CD1d+NKT cells, CD25+T reg cells, CD44hi recirculating memory cells and GL3+ γδ T cells were excluded as dump.
B. Cell proportions of indicated T cell subsets in thymocytes and cell numbers.
C. Representative FACS profiles for CD69+MHCI− (Semi Mature; SM), CD69+MHCI+ (Mature1; M1) or CD69−MHCI+ (Mature2; M2) cells in CD4SP with exclusion of CD1d+NKT cells, CD25+T reg cells, CD44hi recirculating memory cells and GL3+ γδ T cells from 8-week-old mice.
D. Cell proportions of indicated T cell subsets in thymocytes and cell numbers.
E. Representative FACS profiles for Foxp3+CD25+ regulatory T cells and Foxp3–CD25+GITR+ precursor cells from 8-week-old mice.
F. Cell proportions of indicated T cell subsets in thymocytes and cell numbers.
G. Representative FACS profiles for activated CD4 and CD8 T cells in the spleen from 20-week-old control (N = 7), *Ehf*$^{\Delta TEC}$ (N = 4), *Elf3*$^{\Delta TEC}$ (N = 6) and *Ehf*$^{\Delta TEC}$ *Elf3*$^{\Delta TEC}$ (N = 5) female mice.
H. Cell number and percentages of indicated T cell subpopulations. No significant change was found by Tukey's multiple comparisons test.
